# Rate-of-change analysis in palaeoecology revisited: a new approach

**DOI:** 10.1101/2020.12.16.422943

**Authors:** Ondřej Mottl, John-Arvid Grytnes, Alistair W.R. Seddon, Manuel J. Steinbauer, Kuber P. Bhatta, Vivian A. Felde, Suzette G.A. Flantua, H. John B. Birks

## Abstract

Dynamics in the rate of compositional change (rate-of-change; RoC) of biotic or abiotic assemblages preserved in palaeoecological sequences, are thought to reflect changes due to exogenous drivers such as climate and human forcing as well as endogenous factors linked to local dynamics and biotic interactions. However, changes in sedimentation rates and sampling strategies can result in an uneven distribution of time intervals and are known to affect RoC estimates. Furthermore, there has been relatively little exploration of the implications of these challenges in quantifying RoC in palaeoecology.

Here, we introduce *R-Ratepol* – an easy-to-use R package – that provides a robust numerical technique for detecting and summarising RoC patterns in complex multivariate time-ordered stratigraphical sequences. First, we compare the performance of common methods of estimating RoC and detecting periods of high RoC (peak-point) using simulated pollen-stratigraphical data with known patterns of compositional change and temporal resolution. In addition, we propose a new method of binning with a moving window, which shows a more than 5-fold increase in the correct detection of peak-points compared to the more traditional way of using individual levels.

Next, we apply our new methodology to four representative European pollen sequences and show that our approach also performs well in detecting periods of significant compositional change during known onsets of human activity, early land-use transformation, and changes in fire frequency.

Expanding the approach using *R-Ratepol* to open-access paleoecological datasets in global databases, such as Neotoma, will allow future palaeoecological and macroecological studies to quantify major changes in biotic composition or in sets of abiotic variables across broad spatio-temporal scales.

## 1. Introduction

Quantifying spatio-temporal changes in biological composition or diversity is essential for monitoring the current biodiversity crisis and for disentangling underlying drivers such as climate, land-use change, pollution, or introduction of invasive species, as well as intrinsic population dynamics driven by biotic interactions (e.g. Dodd et al., 1994; Dornelas et al., 2014, 2013; Gotelli et al., 2010; Magurran et al., 2010; Monchamp et al., 2018; Morecroft et al., 2009; Richardson et al., 2006; Silvertown et al., 2006; Steinbauer et al., 2018; Wolfe et al., 1987). Therefore, the importance of such ecological long-term observational studies (50-100 years) is increasingly recognised (Dornelas et al., 2014, 2013; Hillebrand et al., 2018; Magurran et al., 2019). Substantial compositional change has been observed during recent decades (e.g. Feeley et al., 2020; Steinbauer et al., 2018), the last few centuries (“The Great Acceleration”; Steffen et al., 2015), and during the Holocene as a result of human impact or in response to regional climate change (Mottl et al., 2021; Seddon et al., 2015; Shuman et al., 2005; Stephens et al., 2019). However, similar rates of substantial change in composition have also been detected on longer (10^2^ to >10^7^ years) geological time scales (Kemp et al., 2015). To understand the impacts of humans, for instance, on ecosystems it is essential to compare temporal changes in species composition and diversity through human history and to investigate whether such changes are unique to the epoch of human impact or whether they precede human-dominated systems (Birks, 1997; Birks et al., 2016; Mottl et al., 2021; Nogué et al., 2021).

Assemblage data of palaeoecological sequences of terrestrial and marine proxies (e.g. pollen, diatoms, chironomids, cladocerans, molluscs, sediment chemical variables, etc.) are an exceptional resource for quantifying spatio-temporal changes in biotic or abiotic composition and diversity beyond the time period of human observations. Rate-of-change (RoC) analysis was introduced into palaeoecology by Jacobson and Grimm (1986) to quantify the rate and the magnitude of compositional change within a Holocene pollen sequence. It was extended by Jacobson et al. (1987) to quantify and compare rates of change within and between sequences (see also Grimm and Jacobson, 1992) in an attempt to identify regional-scale and local-scale patterns in rates of change. RoC analysis estimates compositional change or temporal beta-diversity between adjacent stratigraphical levels and hence between times. Unlike other estimates of beta-diversity, RoC analysis specifically estimates the magnitude of compositional change per *unit time*. Therefore, an essential requirement in RoC analysis is a robust age-depth model for the stratigraphical sequence of interest to obtain the best available estimate of the age of levels within the sequence.

Despite the wide use of RoC analysis in palaeoecology (e.g. Birks and Ammann, 2000; Birks, 1997; Birks and Birks, 2008; Correa-Metrio et al., 2012; Grindean et al., 2019; Laird et al., 1998; Solovieva et al., 2008; Urrego et al., 2009), there has been relatively little exploration of the underlying methodology of RoC analysis and how the various methodologies and sequence properties influence RoC estimates (but see Bennett and Humphry, 1995; Birks, 2012; Lotter et al., 1992). Different approaches to estimate RoC (Bennett and Humphry, 1995; Birks, 2012) involve choices in terms of approach and technique, such as transforming the stratigraphical levels and associated assemblage data (using binning or interpolation to constant time intervals), smoothing the data, and the metric used to estimate the amount of compositional dissimilarity between adjacent levels (Birks, 2012).

The variety of approaches available to RoC analysis can, however, create problems. First, Anderson et al. (2020) stress the sensitivity of RoC to temporal sampling variation within and between records, consequently hampering conclusive comparisons. Second, there has been a lack of standardisation between studies (i.e. incorporating different dissimilarity metrics and temporal procedures) which means RoC outcomes cannot be directly compared between sequences or studies (Birks, 2012). Third, intrinsic properties of an individual sequence, such as its taxonomic richness and the density and distribution of levels within the sequence, can influence the estimated RoC. The end result is that, since the expected patterns of RoC in a sequence are unknown, the most appropriate choices or methods to undertake RoC analysis are also unknown (Bennett and Humphry, 1995; Birks, 2012).

Here, we present and evaluate the performance of *R-Ratepol*, a new R package (R Core Team, 2018), for RoC analysis introducing a standardised and robust method, which incorporates age uncertainties from age-depth modelling as well as standardisation of variable richness, and hence current best practice, for estimating and comparing RoC estimates within and between stratigraphical sequences. First, we present the *R-Ratepol* and its capacities for data commonly used in palaeoecological studies. Second, we compare the performance of various methods of estimating RoC using simulated pollen-stratigraphical data (Blaauw et al., 2010) with known patterns of compositional change and resolution.

In addition, we introduce a method for detecting ‘peak-points’, defined as a rapid change in composition or relative abundances of variables within the sequence, which provides an intuitive, much-needed means to directly compare RoC between sequences and thus facilitates the interpretation of potential drivers of assemblage change on a regional scale. We compare the effectiveness of our new method for the successful detection of peak-points with commonly used methodologies which are also provided in *R-Ratepol*. Finally, we illustrate our findings by estimating RoC values for four pollen palynological sequences with different densities of stratigraphical levels and pollen richness and link the observed patterns to known anthropogenic activities. Our approach can provide important information for future palaeoecological and macroecological studies attempting to quantify, and then attribute, major changes in biotic or abiotic composition across broad spatial areas, and to compare the observed changes in recent decades with changes that occurred within the Holocene or beyond (e.g. Gibson-Reinemer et al., 2015; Mottl et al., 2021).

The term ‘assemblage’ is used through the text to refer to multivariate sets of biotic (pollen, macrofossils, etc.) or abiotic (sediment chemistry, isotope ratios, etc.) variables studied in levels within a stratigraphical sediment sequence.

## 2. Materials and Methods

### 2.1. R-Ratepol

*R-Ratepol* (version 0.6.0) is written as an R package and includes a range of possible settings including a novel method to evaluate RoC in a single stratigraphical sequence using assemblage data and age uncertainties for each level. There are multiple built-in dissimilarity coefficients (DC) for different types of assemblage data, and various levels of data smoothing that can be applied depending on the type and variance of the data. In addition, *R-Ratepol* can use randomisation, accompanied by use of age uncertainties of each level and taxon standardisation to detect RoC patterns in datasets with high data noise or variability (i.e. numerous rapid changes in composition or sedimentation rates).

The computation of RoC in *R-Ratepol* is performed using the following steps (Fig. 1). Detailed descriptions of the underlying methods and relevant formulae are given in the Supplementary Material:

1. Assemblage and age-model data are extracted from the original source and should be compiled together, i.e. depth, age, variable (taxon) 1, variable (taxon) 2, etc.
2. (optional) Smoothing of assemblage data: Each variable within the assemblage data is smoothed using one of five in-built smoothing methods: none, Shepard’s 5-term filter (Davis, 1986; Wilkinson, 2005), moving average, age-weighted average, Grimm’s smoothing (Grimm and Jacobson, 1992).
3. Creation of time bins: A template for all time bins in all window movements is created.
4. A single run (an individual loop) is computed:

a. (optional) Selection of one time series from age uncertainties (see section 2.1.1.2. on randomisation)
b. Subsetting levels in each bin: Here the working units are defined (WUs; see section 2.1.1.1.)
c. (optional) Standardisation of assemblage data in each WU (see section 2.1.1.2. on randomisation)
d. Calculation of RoC between WUs: RoC is calculated as the dissimilarity coefficient (DC) standardised by age differences between WUs. Five in-built dissimilarity coefficients are available: *Euclidean distance*, *standardised Euclidean distance*, *Chord distance*, *Chi-squared coefficient* (Prentice, 1980), *Gower’s distance* (Gower, 1971). The choice of DC depends on the type of assemblage data (see Supplementary Material). In addition, RoC between WUs can be calculated using every consecutive WU, or alternatively, calculation of RoC can be restricted to only directly adjacent WUs. Using the former increases the number of samples for which RoC can be calculated within a sequence, which varies in terms of sample resolution, but may still introduce biases related to the RoC estimation as a result of the varying inter-sample distances.
e. The summary of a single run is produced based on all moving windows
5. Step 4 is repeated multiple times (e.g. 10,000 times).
6. Validation and summary of results from all runs of RoC calculation are produced.
7. (Optional) Data beyond a certain age can be excluded (e.g. 8000 cal yr BP for the pollen sequences in this study)
8. Detection and validation of significant peak-points. There are five in-built methods to detect significant peak-points: *Threshold*, *Linear trend*, *Non-linear trend*, *first derivative* of a generalised additive model (f-deriv GAM; Simpson, 2018), and *Signal-to-Noise Index* (SNI; Kelly et al., 2011).

**Figure 1.**
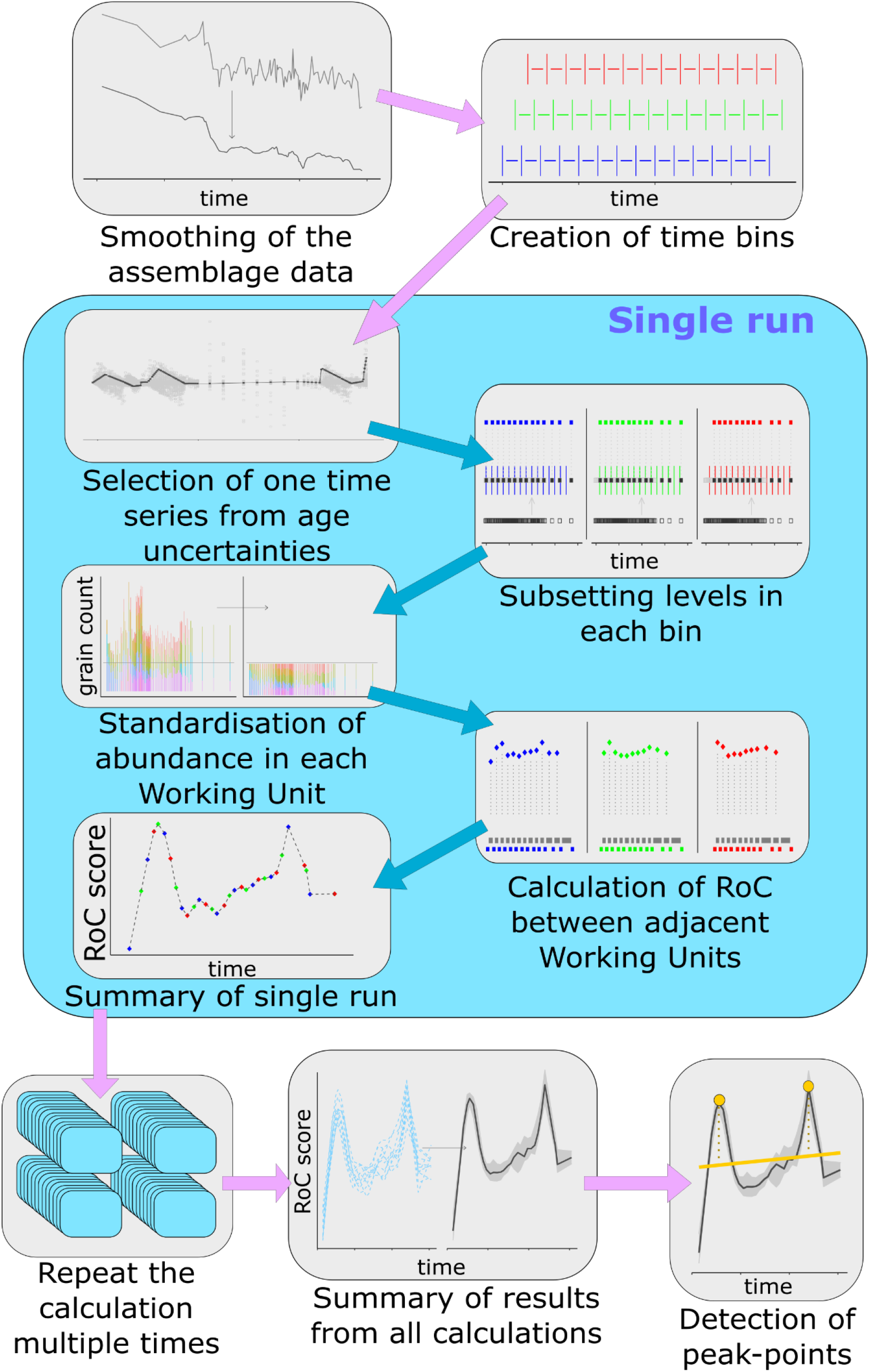
Schematic visualisation of RoC calculation in R-Ratepol using binning with a moving window approach. Methodological retails are provided in section 2.1 and the Supplementary Material.

#### 2.1.1. Specific methodological considerations

##### 2.1.1.1. Selection of working units (WU; Step 3)

RoC is calculated between consecutive Working Units (WU). Traditionally, these WUs represent individual stratigraphical levels. However, changes in sedimentation rates and sampling strategies can result in an uneven temporal distribution of levels within a time sequence, which in turn makes the comparison of RoC between sequences problematic. There are various methods that attempt to minimise such problems. The first is *interpolation of levels* to evenly spaced time intervals, and the use of the interpolated data as WUs. This can lead to a loss of information when the density of levels is high. Second is *binning of levels*: assemblage data are pooled into age brackets of various size (i.e. time bins) and these serve as WUs. Here, the issue is a lower resolution of WUs and their uneven size in terms of total assemblage count (bins with more levels have higher assemblage counts). Third is *selective binning*: like classical binning, bins of selected size are created, but instead of pooling assemblage data together, only one level per time bin is selected as representative of each bin. This results in an even number of WUs in bins with a similar count size in the assemblage. However, the issue of low resolution remains.

Therefore, we propose a new method of *binning with a moving window*, which is a compromise between using individual levels and selective binning. This method follows a simple sequence: time bins are created, levels are selected as in selective binning, and RoC between bins is calculated. However, the brackets of the time bin (window) are then moved forward by a selected amount of time (Z), levels are selected again (subset into bins), and RoC calculated for the new set of WUs. This is repeated X times (where X is the bin size divided by Z) while retaining all the results.

*R-Ratepol* currently provides several options for selecting WU, namely as individual levels, selective binning of levels, and our new method of binning with a moving window.

##### 2.1.1.2. Randomisations

Due to the inherent statistical errors in uncertainties in the age estimates from age-depth and the assemblage datasets (e.g. pollen counts in each level; Birks and Gordon, 1985), *R-Ratepol* can be run several times and the results summarised (Steps 5-6). Therefore, two optional settings are available by using age uncertainties and assemblage data standardisation.

###### Age uncertainties (Step 4a)

For each run, a single age sequence from the age uncertainties is randomly selected. The calculation between two consecutive WUs (i.e. one working-unit combination) results in a *RoC score* and a *time position* (which is calculated as the mean age position of the two WUs). However, due to random sampling of the age sequence, each WU combination will result in multiple RoC values. The final RoC value for a single WU combination is calculated as the median of the scores from all randomisations. In addition, the 95^th^ quantile from all randomisations is calculated as an error estimate.

###### Data standardisation (Step 4b)

Variables (taxa) in the assemblage dataset can be standardised to a certain count (e.g. number of pollen grains in each WU) by rarefaction. Random sampling without replacement is used to draw a selected number of individuals from each WU (e.g. 150 pollen grains).

#### 2.1.2. Detection of peak-points in RoC sequence (Step 8)

A rapid change in composition or relative abundances of variables within the sequence can provide a means of comparing RoC between sequences and interpreting the potential drivers of assemblage change. To detect such significant peak-points of RoC scores in each sequence, each point is tested to see if it represents a significant increase in RoC values. There are various ways to detect peak-points in a time series and *R-Ratepol* is able to detect peak-points using five methods:

1. Threshold: Each point in the RoC sequence is compared to a median of all RoC scores from the whole sequence (i.e. *threshold value*). The ROC value for a point is considered significant if the 95^th^ quantile of the RoC scores from all calculations is higher than the *threshold value*.
2. Linear trend: A linear model is fitted between the RoC values and their ages. Differences between the model and each point are calculated (residuals). The standard deviation (SD) is calculated from all the residuals. A peak is considered significant if it is 1.5 SD higher than the model (this value can be selected by the user).
3. Non-linear trend: A conservative generalised additive model (GAM) is fitted through the RoC scores and their ages (GAM = RoC ~ s(age, *k* = 3) using the *mgcv* package (Wood, 2011). The distance between each point and the fitted value is calculated (residuals). The standard deviation (SD) is calculated from all the residuals. A peak is considered significant if it is 1.5 SD higher than the model (this value can be selected by the user).
4. F-deriv GAM: A smooth GAM model is fitted to the RoC scores and their ages (GAM = RoC ~ s(age). The first derivative as well as continuous confidence intervals are calculated from the model using the *gratia* package (Simpson, 2019). A peak is considered significant if the confidence intervals of the first derivative differ from 0 (for more information see Simpson, 2018).
5. Signal-to-noise (SNI) method: We adapted SNI from Kelly et al. (2011), which was developed to detect changes in charcoal stratigraphical records. SNI is calculated for the whole RoC sequence and a peak-point is considered significant if it has an SNI value higher than 3.

### 2.2. Testing the successful detection of peak-points using simulated data

In order to compare the performance of various methods of estimating RoC, we used simulated pollen-stratigraphical data (Blaauw et al., 2010) with known patterns of compositional change and temporal resolution to test the success of peak-point detection in the expected time period of compositional change. In addition, we used a generalised linear mixed modelling approach to evaluate the effects of the different *R-Ratepol* parameters. Finally, we performed a series of sensitivity tests comparing the binning with the moving window approach to other methods and to test the robustness of the RoC estimation.

#### 2.2.1. Data simulation

We simulated pollen datasets following Blaauw *et al.* (2010) based on generating pollen-assemblage data in response to known changes in environmental conditions and compositional properties (Fig. S1). We applied the following process of data generation: (i) the density of levels from a European pollen sequence, Glendalough (Sequence A), was used as a template for the number of levels and the corresponding ages for each level; (ii) Pollen data were created as abundance data with a total of 300 pollen grains in each level; (iii) We then adjusted the total values of each simulated taxon so that the pollen assemblage resembled a log-scale rank distribution; (iv) We added jitter (random noise) to each of the pollen taxa (function *jitter*, factor = 1.5). All changes were made in order to make the simulated data more similar to data from a real-life study.

Two additional settings were altered during the simulation of the datasets: (i) richness – *low richness* (LR) datasets contain 5 pollen taxa, *high richness* (HR) datasets contain 50 pollen taxa; and (ii) position of change in external environmental properties – *late* (L) has a sudden increase of environmental properties at 2000 calibrated years before present (1950 CE: cal yr BP) and a decrease at 3000 cal yr BP, *early* (E) has a sudden increase at 5500 and a decrease at 6500 cal yr BP (Fig. S1). This results in two abrupt changes in pollen composition in each dataset. Note that the different timings of the change in environmental properties (hereafter called ‘focal period’) were selected to illustrate the effects of level density (datasets with the *early* change have a low density of levels within the focal period and datasets with a *late* change have a high density). The combination of these two settings results in four types of simulated datasets: *low richness – late change* (LR-L)*, low richness – early change* (LR-E)*, high richness – late change* (HR-L), and *high richness* – *early change* (HR-E).

#### 2.2.2. Using the simulated datasets to test analytical performance of RoC methods with different settings

To investigate the differences between the various settings in RoC methods, we simulated 100 datasets for each dataset type (LR-L, LR-E, HR-L, HR-E; see previous section), and compared the success rate of detecting peak-points in the expected time period for all combinations of (i) WU selection, (ii) smoothing method, (iii) dissimilarity coefficient, and (iii) peak-point detection.

We tested three types of WU selection (individual stratigraphical levels, selective binning, and binning with a moving window; RoC estimation was restricted to only directly adjacent WUs in all calculations), five types of smoothing methods (none, Shepard’s 5-term filter, moving average, age-weighted average, Grimm’s smoothing), two types of dissimilarity coefficients appropriate for closed compositional data (Chord distance, Chi-squared coefficient), and five types of peak-point detection (Threshold, Linear trend, Non-linear trend, f-deriv GAM, SNI). Of these, all the methods are well established, with the exception of i) binning with a moving window, ii) threshold, iii) linear trend, and iv) non-linear trend, which are new methods developed here. In addition, we did not include Euclidean and standardised Euclidean dissimilarity coefficients in the calculation, as they are not appropriate for percentage pollen data (Prentice, 1980).

For each simulated dataset, we calculated RoC using combinations of all WU selections, data smoothing, dissimilarity coefficient, and peak-point detection method (150 different settings in total). *R-Ratepol* has been set up with standardisation of pollen assemblage data to 150 pollen grains in each WU and 100 randomisations per calculation. The number of randomisations is not higher due to high computational demands (60,000 RoC calculations, each with 100 randomisations). Randomisations are in place only for taxon standardisation as no age uncertainties have been used in this example. We then extracted (i) the number of WUs where *R-Ratepol* detected a significant peak-point during the focal period (± 500 yr), and (ii) the number of levels where *R-Ratepol* detected a peak-point during a time of no expected change (‘false positives’). We transformed these numbers of peak-points into ratios of points detected to total points in the area (inside or outside focal period; see Figs S2 and S3 for examples of differences in successful peak-point detection between different WU selection method and peak-point detection methods).

##### Statistical tests to evaluate RoC options

First, we tested if the various WU selection methods affect the successful detection of peak-points within the focal period. We pooled all RoC calculations from all dataset types and created generalised linear mixed models using the Template Model Builder (Brooks et al., 2017) glmmTMB _*success*–*method*_ with the ratio of WU marked as peak-points to all WUs in the focal period (*R*_*success*–*method*_) as the dependent variable with a beta error distribution. Independent variables were: WU (3 level factor), peak-point detection method (5 level factor), dataset type (4 level factor), and all their interactions. Individual dataset ID and RoC setting (factor combining smoothing method and DC; 10 levels) were selected as the random factors. We then used the *dredge* function from the *MuMIn* package (Barton, 2020) to fit models with all possible combinations of predictors (with a constraint that dataset type must be present), and ordered the models by parsimony (assessed by the Akaike information criterion; *AIC*_*C*_). We only selected the best model if Δ*AIC*_*C*_ (i.e. *delta* in the *MuMIn* package) is < 2. In the case where multiple models had similar parsimony, we selected the best model using the *compare_performance* function from the *performance* package (Ludecke et al., 2020). We then used the *emmens* package (Russell, 2020) to obtain estimated marginal means and the 95^th^ quantile of the independent variables from the final model. Similarly, we built glmmTMB _*FalsePositive*–*method*_ using the ratio of WUs marked as peak-points to all WUs outside the focal period (*R*_*FalsePositive,*–*method*_).

Next, for a more detailed exploration of different settings in *R-Ratepol*, we divided the data to only include the best performing WU selection and peak-point detection combination (i.e. binning with a moving window and non-linear trend peak-point detection, see Results) and created glmmTMB _*success*–*detail*_ and glmmTMB _*FalsePositive*–*detail*_ with *R*_*success*–*detail*_ and *R*_*FalsePositive*–*detail*_ as the dependent variables, respectively. Both models were fitted with a beta error distribution and the independent variables are data-smoothing type (5 level factor), dissimilarity coefficient (2 level factor), position of environmental change (i.e. density of levels; 2 level factor), richness of dataset (2 level factor), and all their interactions. Individual dataset ID was selected as a random factor. As before, the *dredge* and *compare_performance* functions were used to select the best model and reduce unnecessary predictors.

#### 2.2.3. Sensitivity analyses

We conducted a series of analyses to compare methods of WU selection (individual stratigraphical levels, selective binning, and binning with a moving window) and to test the robustness of results using scenarios of: (i) changing size of time bin, (ii) various level resolutions, (iii) hiatus between levels, and (iv) missing levels from the top of the sequence. For this purpose, we used a European pollen sequence, Glendalough (Sequence A, see section 2.3.), with its assemblage data smoothed using age-weighted average, because it represents a sequence with one of the main challenges represented in RoC estimation, i.e. varying sample resolution across the record. First, to the test of varying bin sizes (i), we used binning with a moving window and estimated RoC using time bin sizes ranging from 100 to 3000 yr. Second, to test the effect of levels resolution on our RoC estimation (ii), we created a series of datasets with varying level resolution, starting from the full dataset to a dataset with an inter-level distance of at least 500 yr. We then estimated RoC for each of those datasets using three methods of WU selection (individual stratigraphical levels, selective binning, and binning with a moving window). Since the length of the time bin has to be larger than the inter-sample distance to estimate RoC, we selected a time bin of 1000 yr. Third, for the test of the effect of a hiatus in the levels (iii), we created a series of datasets and manually implemented hiatuses of various length starting from full dataset to hiatus of 2000 yr. All hiatuses began at 3000 cal yr BP. The size of the time bin was selected at 500 yr. Finally, for the test of missing levels from the top of the sequences (iv), we followed similar methodology as in (iii) but levels were removed starting from the top of the sequence. All RoC scores are expressed as dissimilarity per 500 yr. The option to only calculate RoC between non-adjacent units was turned off for these sensitivity analyses. Note that repeating these sensitivity tests with this option turned on gave overall similar results.

### 2.3. Examples of rate-of-change results using empirical palynological data

After assessing the differences between the individual settings of *R-Ratepol* based on the results from the simulated datasets, we examined the RoC estimates for real palynological data representing different pollen richness and composition, and different density of levels. We obtained the pollen data from the Neotoma database (Williams et al., 2018) using the *Neotoma R package* (Goring et al., 2015). We chose four European sequences (A–D; Fig 2).

**Figure 2.**
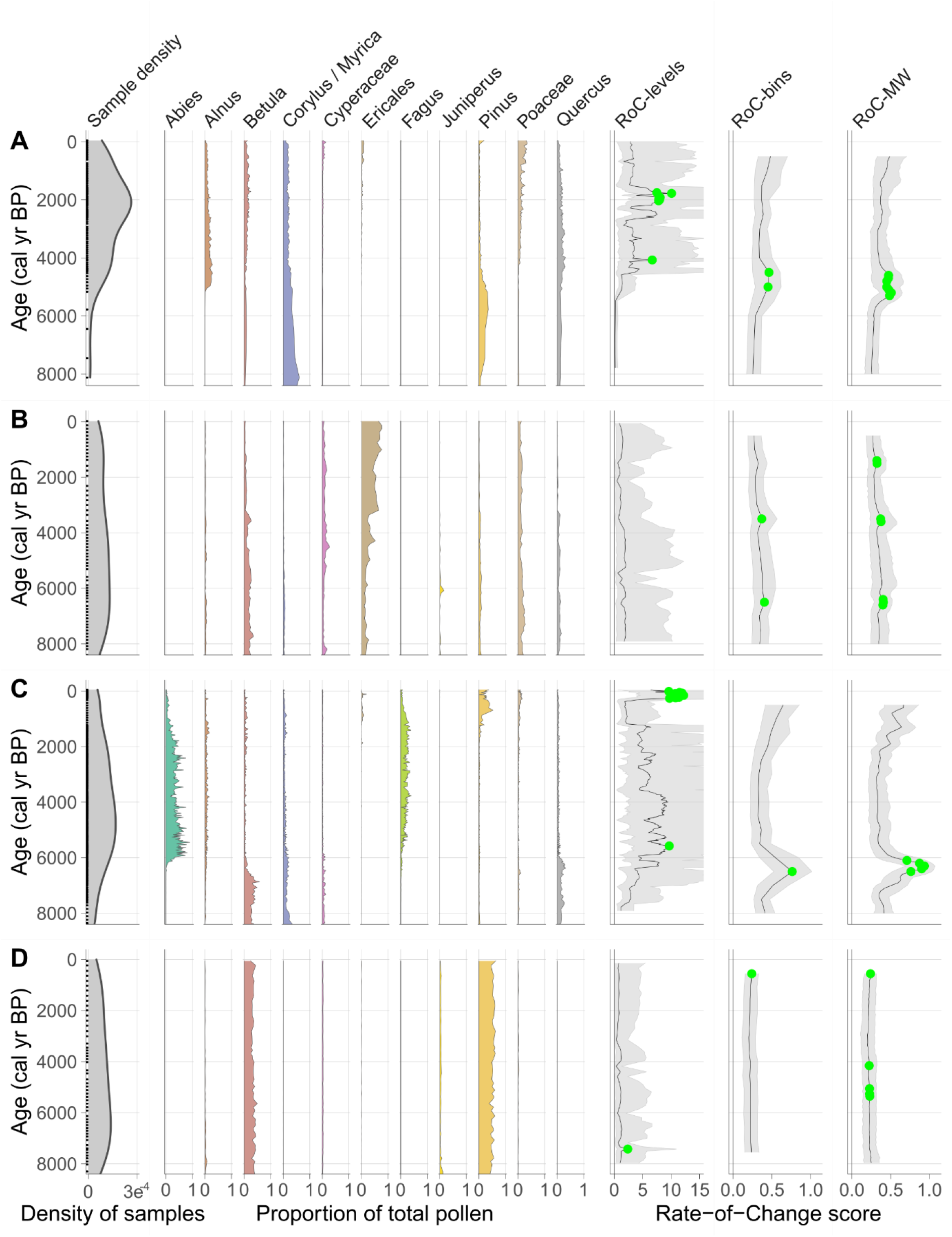
Summary of four sequences used in this study. Glendalough (A), Dallican Water (B), Steerenmoos (C), and Alanen Laanijärvi (D). For each sequence, three different plots are shown. First plot (left) shows the density of levels within each sequence for the last 8000 cal yr BP. Second plot (middle) shows pollen stratigraphies for the most common taxa in the sequences. Values are presented as proportions of pollen grains in each level. Last plot (right) shows the rate-of-change score calculated with *R-Ratepol* for three different methods for WU selection (RoC score is represented as dissimilarity per 500 yr): RoC-levels = use of individual levels, RoC-binning = binning, RoC-MW = binning with moving window. Age-weighted average smoothing and Chi-squared coefficient were selected for smoothing and dissimilarity measure, respectively. Green points indicate significant peak-points detected using the non-linear trend method. For a detailed explanation of the methods, see Materials and Methods, and Supplementary Material.

In each sequence, taxa were harmonised to the taxonomically highest pollen morphotype (Level = MHVar2) using the pollen harmonisation table in Giesecke et al. (2019). To develop age-depth models, we used the pre-selected radiometric control points provided in Giesecke et al. (2013) and calibrated the radiocarbon dates using the IntCal13 Northern Hemisphere calibration curve (Reimer et al., 2013). For each sequence, we constructed an age-depth model using the *Bchron* R package (Haslett and Parnell, 2008) to generate 1000 possible age estimates for all sample levels at the sampling resolution of the original pollen sequences. We used these 1000 draws to build posterior estimates of age uncertainty. We calculated the median-age estimate for each sample level to obtain the default age used in subsequent analyses.

In each sequence, we excluded all levels that contained less than 150 pollen counts of terrestrial taxa, and all levels beyond a 3000-year extrapolation of the oldest chronological control point. In addition, we excluded all levels with an age older than 8500 cal yr BP to ensure a focus on the period with detectable human impact.

**Sequence A** is from Glendalough, Ireland (53°00’10.0”N 6°22’09.0”W; Neotoma dataset id = 17334; Haslett et al., 2006). It has a very uneven distribution of levels (N = 102) with the highest density around 2000 cal yr BP (Fig. 2A). The sequence contains 80 pollen taxa with abundant *Corylus*, *Quercus, Alnus, Betula*, and Poaceae. Cyperaceae, Poaceae, and Ericales all increase in the last 1000 cal yr BP, preceded by a period with high values of *Quercus*, *Betula*, and *Alnus* until ~ 5000 cal yr BP. Before 5000 cal yr BP the sequence has large variations in *Pinus*, an increase of *Alnus*, and a large decrease in *Corylus*. **Sequence B** is from Dallican Water, Scotland (60°23’14.5”N 1°05’47.3”W; Neotoma dataset id = 4012; Bennett et al., 1992). It has a relatively even distribution of levels (N = 63; Fig. 2B) and contains 50 pollen taxa with Ericales, *Betula*, and Poaceae being the most abundant. The pollen record shows sudden increases of Ericales after 4000 cal yr BP. **Sequence C** is from Steerenmoos, Germany (47°48’20.0”N 8°12’01.6”E; Neotoma dataset id = 40951; Rösch, 2000). It is an example of a very detailed sequence, but with an uneven distribution of levels (N = 273; Fig. 2C). The sequence contains 103 pollen taxa with *Abies*, *Fagus*, *Corylus, Betula*, and *Quercus* being the most abundant. As with Sequence A, the pollen stratigraphy can be separated into three major parts: i) a recent period until 1000 cal yr BP typified by high values of *Pinus*; ii) 2000–6000 cal yr BP characterised by *Abies* and *Fagus*; and iii) 6000 cal yr BP to the base with abundant *Betula*, *Corylus*, and *Quercus*. **Sequence D** is from Alanen Laanijärvi, a boreal-forest lake in Swedish Lapland (67°58’N 20°29’W; Neotoma dataset id = 45314; Heinrichs et al., 2005). It is an example of a sequence with 54 levels across the time of interest, containing 44 pollen taxa with a high abundance of *Pinus* and *Betula* throughout the whole sequence (Fig. 2D).

We calculated RoC scores for the four selected European sequences (A-D) using all three methods of WU selection with age-weighted average smoothing and Chi-squared coefficient as the dissimilarity coefficient. A non-linear trend peak-point detection was used in each sequence, the number of randomisations was set to 1,000, and the size of time bin was selected as 500 yr (see Fig. S4 for an example of changes of RoC values with the size of time bin). RoC estimation was restricted to only directly adjacent WUs in all calculations

In addition, to provide examples of the differences between different settings of *R-Ratepol*, we explored data from sequence A using binning with the moving window method of WU selection and calculated RoC scores for all combinations of five smoothing methods and two dissimilarity coefficients (Chi-squared, Chord distance).

## 3. Results

### 3.1. Comparison of success rates in peak-point detection for simulated datasets

Successful detection of peak-points (*R*_*success*–*method*_) and the number of incorrectly (false positives) detected peak-points (*R*_*FalsePositive*–*method*_) are both significantly affected by WU selection and the method of peak-point detection (Fig. 3; see Table S1 and Table S2 for the *AIC*_*C*_ of all models). For detecting peak-points successfully within the focal period, our approach of binning with a moving window results in an overall better performance in successful peak-point detection than the use of individual levels or binning (effect of successful detection increases by more than 5-fold and 3-fold, respectively; Fig. 3A). Simulations show that the non-linear trend method performs best in detecting peak-points within the focal period (16% higher than the second-best, the linear trend method; Fig. 3B), and detects a similar amount of false positives compared to the other peak-point detection methods (with the exception of the first derivative of a GAM (GAM first deriv), which detects 76% more false positives).

**Figure 3.**
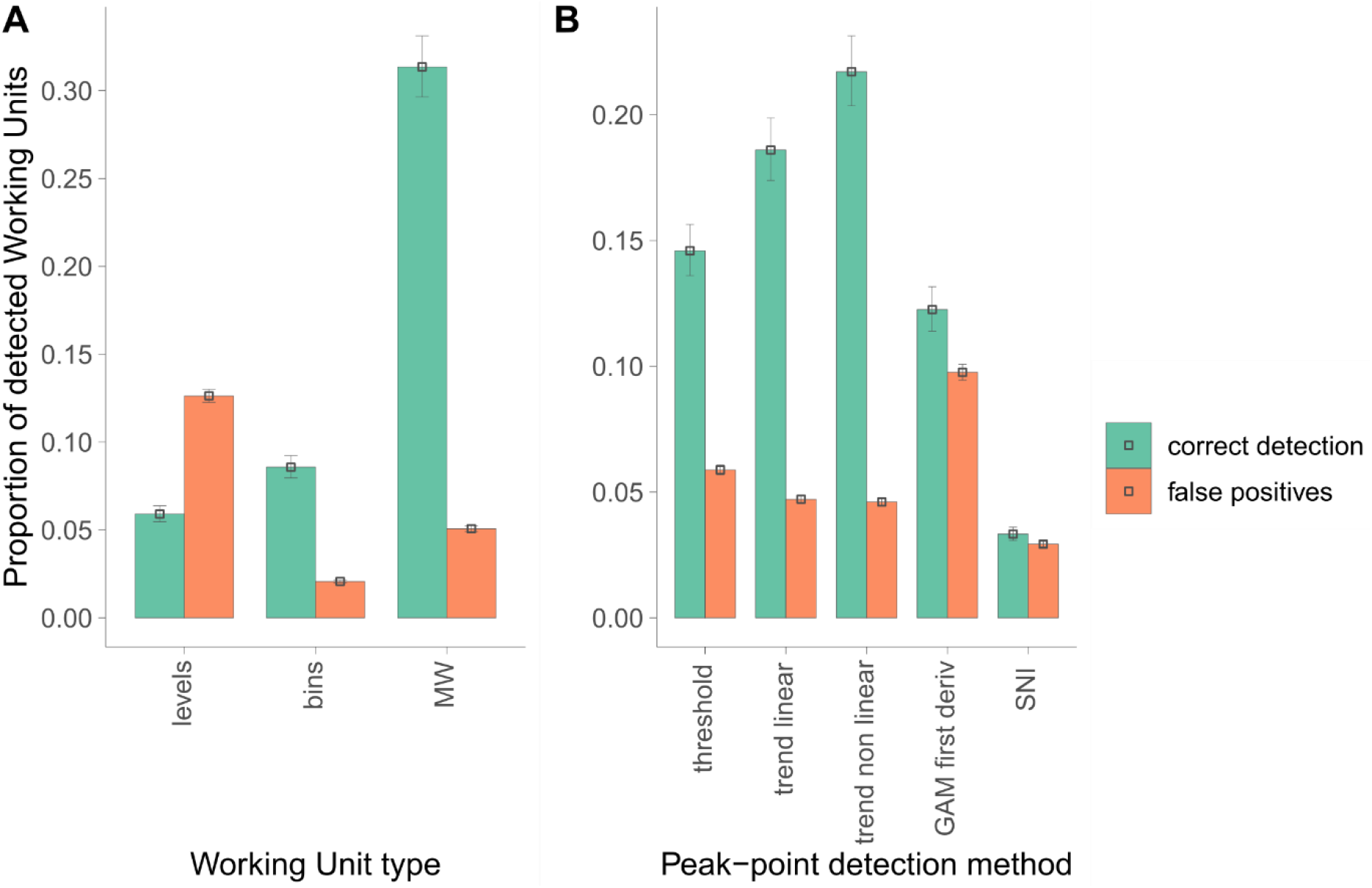
Comparison of the correct detection of peak-points in simulated datasets. A) Comparison of estimated marginal means of peak-point detection rates between different WU selections. Working unit selection: levels = use of subsequent levels, bins = selective binning, MW = binning with moving window. B) Comparison of estimated marginal means of peak-point detection rates between different peak-point detection methods. GAM first deriv = first derivative of a GAM; SNI = signal-to-noise index.

While using only the binning with the moving-window approach and the non-linear trend method for peak-point detection, the successful detection of peak-points (*R*_*success*–*detail*_) and the number of false positives (*R*_*FalsePositive*–*detail*_) are both influenced by the density of levels, type of dissimilarity coefficient, and data-smoothing type (Fig. 4; see Tables S3, S4, S5, and S6 for the best selected models). Using Chi-squared coefficient as the dissimilarity measure results in a similar number of correctly detected peak-points as Chord distance does on average (Fig. 4; Fig. S5A). The level of smoothing has, on average, a similar effect on the successful detection of peak-points (Fig. S5B). However, due to the interaction between smoothing and the position of environmental change, data smoothed using an age-weighted average and Grimm’s smoothing were the most successful approaches (Fig. 4). The number of false positives is similar among the smooth methods. The datasets with an assemblage change which occurred during a time period characterised by a high density of levels (i.e. late environmental change) show, on average, a 24% lower success of peak-point detection and a 100-fold higher false positive detection than datasets with a change at the time of a low density of levels (i.e. environmental change early in the sequence; Fig. S4C). Richness of pollen does not show a significant effect on the detection of peak-points (note that assemblage data standardisation was used for all computations).

**Figure 4.**
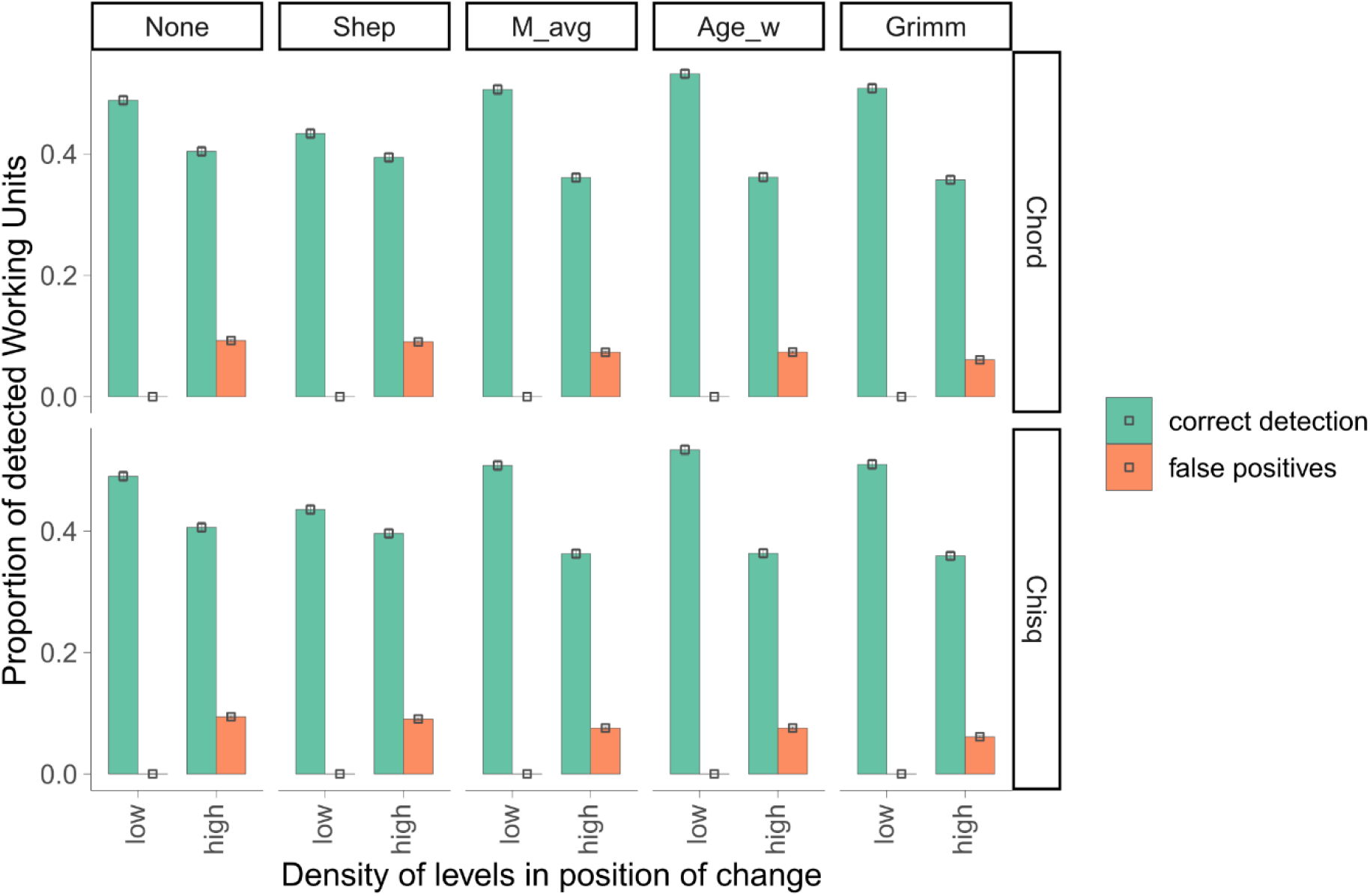
Comparison of peak-point detection rates using binning with a moving window and non-linear peak detection method. Smoothing methods: None = data without smoothing, Shep = Shepard’s 5-term filter, M_avg = moving average, Age_w = age-weighted average, Grimm = Grimm’s smoothing. Dissimilarity coefficients: Chord = Chord distance, Chisq = Chi-squared coefficient. For a detailed explanation of the methods, see Supplementary Material.

### 3.2. Examples of rate-of-change results using real palynological data

The selection of WUs affects not only the overall shape of the RoC curve in all sequences but also the timing of the period of significant increase in RoC (Fig. 2). Binning with a moving window detects significant periods of increased RoC scores at sequence A (Glendalough) for 5300–4600 cal yr BP; sequence B (Dallican Water) for 1500-1400, 3600–3500, and 6600-6400 cal yr BP; sequence C (Steerenmoos) for 6500–6100 cal yr BP; and sequence D (Alanen Laanijärvi) for 550, 4100, and 5300-5000 cal yr BP (Fig. 2). The binning method shows relatively similar patterns and position of peak-points (except for sequence D). Using individual levels as WUs (Fig. 2. ‘RoC-levels’) results in very different positions of peak-points, a different scale of RoC values, and a different shape of the RoC curve with wide error intervals.

### 3.3 Sensitivity tests

#### 3.3.1. Effects of time-bin size (Fig. S4)

Changing the time-bin size overall affects the RoC score, with an increasing mean RoC with decreasing size of time bins. In general, except for the extremes (bin size of 2000 years), the same overall features of the pollen-assemblage changes are expressed by the different bin sizes, although the overall RoC patterns shift through time as the mean-age for the first sample increases with increasing bin size. At the smallest bin size (100 years), the changes in the top of the record are amplified. Note that applying the same sensitivity test to an RoC estimation, where non-adjacent WU were used to estimate RoC also reveals a trend in the overall RoC magnitude. Nevertheless, the overall patterns in RoC using this option are similar, especially when rescaled to the same unit mean and variance. Overall, these results indicate the importance of using the same binning size for all sequences when comparing RoCs between multiple sequences.

#### 3.3.2 Effects of changing the level resolution (Fig. 5)

Changing the effect of level resolution has the largest effect on RoC estimation if no binning procedure is applied. As sample resolution reduces, RoC scores estimated without any binning result in large differences in RoC estimations across the record. For example, the high RoC scores between 2000 and 3000 yr cal BP (mainly the result of reduced inter-levels temporal distance) are gradually reduced relative to other periods in the record. In contrast, RoC estimates using binning with a moving window and the binning approaches result in relatively stable RoC curves irrespective of level resolution, although a number of peak-points are detected in the top of the sequence in the moving-window approach, whilst the peak-points are not detected at the top of the sequence when the standard binning approach is used. Here the patterns in RoC from the two binning approaches generally reflect the overall patterns of assemblage change as observed in the sediment records (i.e. the expansion of *Alnus* and corresponding decrease in *Pinus* pollen at 5000 years, and an increase in the relative abundances of Poaceae, Cyperaceae, and Ericales in the last 1000 years).

**Figure 5.**
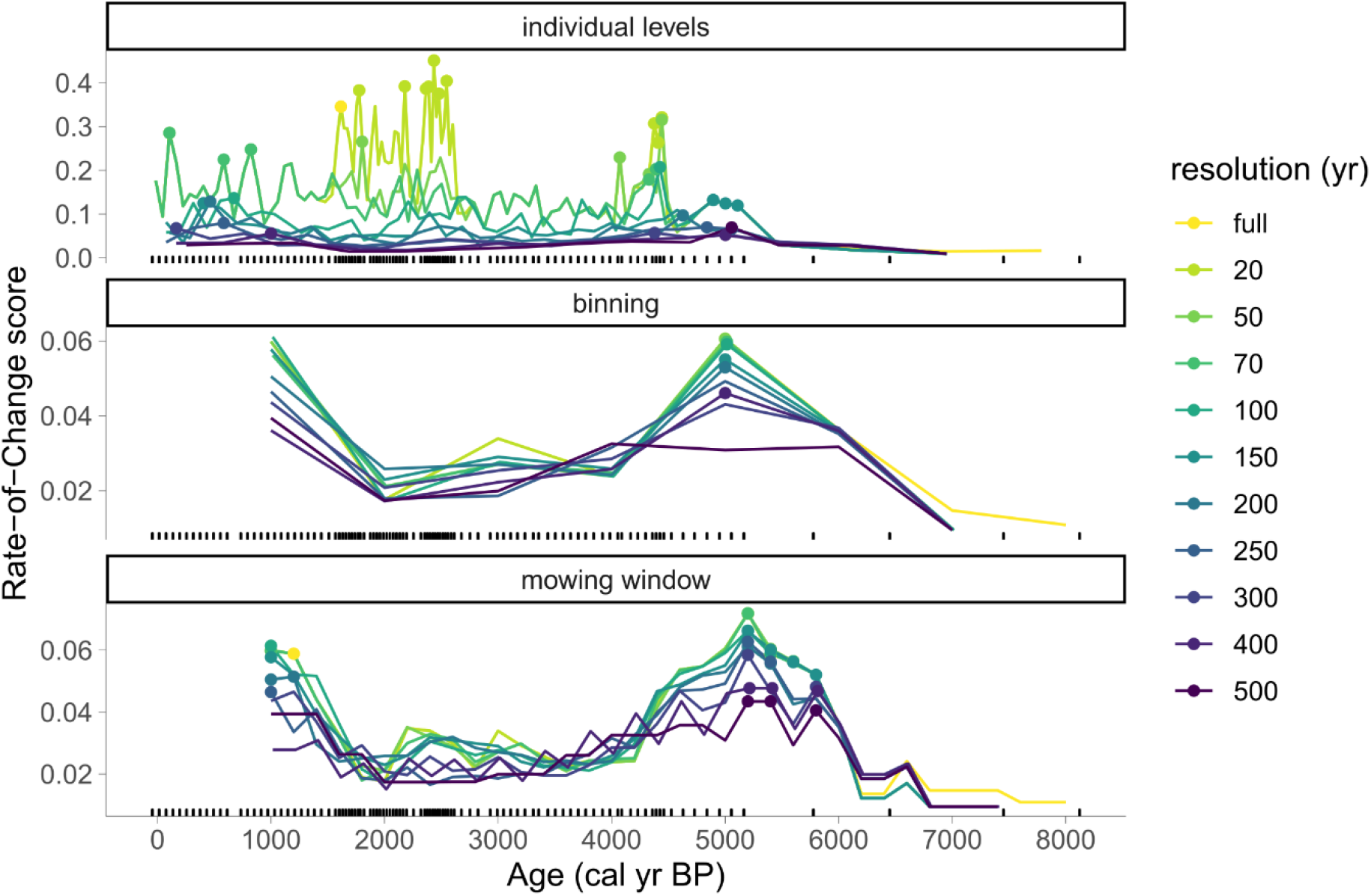
Sensitivity analyses showing RoC scores for datasets with different level resolutions, estimated using three different WU selection methods (individual stratigraphical levels, selective binning, and binning with a moving window). A series of datasets with various level resolution were created from Sequence A (Glendalough). Age-weighted average and Chi-squared coefficient were selected for smoothing and dissimilarity. The time bin for binning and binning with the moving window was set as 1000 yr. Black ticks on the x-axis indicate the positions of the stratigraphical levels (full dataset). RoC score is represented as dissimilarity per 500 yr. Note that the high rates of change detected in RoC in the individual-level approach between 2000 and 3000 years (top panel) are likely the result of the changing sedimentation rates in the core. The binning approaches effectively smooth these effects.

#### 3.3.3 Effects of hiatuses (Figures S6 and S7)

The effects of introducing a hiatus in levels at 2000 years has no effect on the RoC estimation for parts of the sequence, where hiatuses are not present, for both the moving window and binning approaches. Similarly, binning approaches reveal consistent results when dealing with hiatuses found at the beginning of the sequence: For both moving window and binning approaches, the RoC estimates are consistent irrespective to the number of levels removed from the beginning of the sequence.

## 4. Discussion

Based on the results of our simulations, we have shown that the choice of methods and parameters is critical in the analysis of rate-of-change. Comparison of methods using simulated datasets shows that selection of WUs, peak-point detection method, dissimilarity coefficient, smoothing of the data, and the density of the levels in the period of environmental change significantly influence the detection of sudden changes in pollen stratigraphical sequences. Our proposed method of binning with a moving window in combination with a non-linear trend peak-point detection method improves such detection of an increased RoC. The method shows more than a 5-fold increase in the effect of correct detection of peak-points compared to the more traditional approach of selection by individual levels. Peak-point detection using a non-linear trend provides a reasonable compromise between statistical error Type I (linear trend and threshold method) and Type II (SNI method). In addition, binning with a moving window can effectively detect peak-points in a scenario with varying level resolutions (Fig. 5) and produces stable results even when a level hiatus is present in the sequence (Fig. S6-S7).

There is little difference between using Chi-squared coefficient or Chord’s distance as the dissimilarity coefficient for pollen compositional data, and we therefore recommend using both for palaeoecological data sequences that are expressed as percentages or proportions and that contain many zero values. However, *R-Ratepol* is created as modular tool and can be used for many different palaeo-proxies present in sedimentary sequences. Euclidean distance (see Supplementary Material) is appropriate for absolute assemblage data such as pollen concentrations or pollen accumulation rates (e.g. Peglar, 1993; Peglar and Birks, 1993). Such data may require a log-transformation prior to calculating the Euclidean distance between levels or WUs. Standardised Euclidean distance (see Supplementary Material) is appropriate when the variables in an assemblage are expressed in different units, such as geochemical, isotope, and sediment variables (e.g. Bakke et al., 2009; Birks and Birks, 2006; Nesje et al., 2014). Gower’s (1971) distance is appropriate for assemblage data consisting of different numerical types (continuous quantitative, qualitive, binary) such as plant macrofossil data (e.g. Birks, 2014). For more detailed information about the selection of different dissimilarity coefficients in various multivariate data, see Prentice (1980) and Legendre and Legendre (2012).

Recommendation of a specific smoothing technique of data is less straightforward. Of course, there is a trade-off between the potential loss of information by smoothing the data and the incorrect attribution of noise (for example from sampling error) as signal. Our simulation experiments show that while using only binning with a moving-window approach and the non-linear trend method for peak-point detection, data with age-weighted average and Grimm’s smoothing perform the best (Fig. 3). However, detailed exploration of the RoC pattern in a real pollen sequence (sequence A; Fig. S8) shows that all smoothing methods result in a similar RoC curve and position. Therefore, there is a need for critical evaluation in future studies to assess the risks of using data smoothing, depending on the density of levels and the research questions of interest.

A multitude of appropriate data-smoothing approaches (Wilkinson, 2005), dissimilarity coefficients (e.g. Legendre and Legendre, 2012; Prentice, 1980), techniques for creating WUs, and methods of peak-point detection (e.g. Simpson, 2018) are potentially available. Of the two selected dissimilarity methods and five smoothing algorithms, all commonly used in palaeoecology, our model results show that datasets with a high density of levels tend to have a lower chance of successfully detecting peak-points when using binning with a moving window and the non-linear trend method for peak-point detection (Fig. S3C). However, this is probably caused by our method of success assessment. Binning with a moving window results in the correct detection of peak-points in the expected period but there is also a high number of points not identified as peak-points, due to the higher resolution of the curves and the method being able to detect the peak-point increases more precisely (Fig. S2). Therefore, we do not see this as a disadvantage of the method. Traditional methods of using individual levels are more drastically affected by the sample density, as more levels in the sequence result in a higher chance of a very high RoC score only due to a higher level density. This pattern can be observed in sequence A (Glendalough) and C (Steerenmoos), where peak-points were detected in periods with an increased density of levels. Anderson et al. (2020) reached similar conclusions (high density of levels is recommended to successfully detecting RoC patterns) in a regional-scale synthesis of fossil pollen data from California. Here these authors suggest maintaining a consistent temporal spacing within records, and the use of probabilistic models explicitly incorporating age-model uncertainties to increase the precision of age estimates of each level. We show that our method of binning with a moving window (which does not require a consistent temporal spacing) yields even better results than the traditional use of individual levels or binning, and has an advantage here as age-model uncertainties can be incorporated. A sensitivity analyses on sequence A (Fig. 5) confirms that the method of binning with a moving mowing is better at dealing with an uneven level density, even with decreasing resolution. This method is able to detect peak-points in sequences with a very low level resolution and does not show false positives where there is a very high level resolution (Fig. 5). Nevertheless, it remains important to standardise the length of working units if working across multiple cores (Figure S4).

RoC analysis has several applications in palaeoecological research. It is a useful numerical tool to detect patterns in stratigraphical data that cannot readily be seen by visual inspection (see Jacobson et al., 1987 for an example at Gould Pond, Maine). It is also useful to compare the rates of change in different proxies (pollen, diatoms, chironomids, etc.) studied in the same stratigraphical sequence (e.g. Birks and Ammann, 2000; Szabó et al., 2020). Moreover, it is most useful when RoC results are compared from several sequences (e.g. Grimm and Jacobson, 1992; Jacobson et al., 1987; Mottl et al., 2021). Consistent patterns in RoC peaks in several sequences may potentially indicate responses to exogenous drivers such as regional climate change, pathogenic attacks, or widespread human activity (Mottl et al., 2021). Ecologists are recognising the importance of quantifying RoC of environmental drivers such as temperature, toxins, and salinity on ecosystems (Pinek et al., 2020) and are developing new and powerful numerical tools for space–time analysis of community compositional data over time intervals of decades (e.g. Legendre and Gauthier, 2014). RoC peaks unique to individual sequences may, in contrast, reflect local endogenous factors such as sediment reworking or natural disturbance.

A connection between RoC and local factors can be seen in the sequences used in this study (Fig. 2). Sequence A (Glendalough) shows significant peak-points at the time of the onset of human activity and the expansion of grassland and heath (Haslett et al., 2006). Sequence B (Dallican Water) has significant peak-points at times when major changes in land-use occurred as the island was abandoned by humans and then reinhabited 800 years later (Bennett et al., 1992). Sequence C (Steerenmoos) has several significant peak-points at the time of the expansion of *Abies* pollen in the late Neolithic period and with frequent fires (Rösch, 2000). Sequence D (Alanen Laanijärvi) has significant peak-points in the very recent period, possibly associated with changes in organic deposition, fertilisation, and timber harvesting (Heinrichs et al., 2005). Here we can assume that our approach of binning with a moving window is closest to the expected pattern, further supported by the relatively high, successful detection of peak-points in the simulated datasets (Fig. 3, Fig. 5). In contrast, the traditional use of individual levels fails to detect the periods of significant change, or falsely assigns peak-points to other, non-relevant periods, or a combination of both (Fig 2, Fig. 5). Some of these errors occur, as expected, with a low density of levels. These sequences and the RoC outcomes exemplify the usefulness of quantifying rates of compositional change estimated through time.

In summary, we have developed a framework for the robust estimation of rate-of-change in stratigraphical time-ordered sequences of terrestrial and marine palaeoecological datasets. Our overall framework consists of four major parts:

1. Establishing a robust age-depth model for the stratigraphical sequence of interest with age uncertainties for all individual levels.
2. Selecting consecutive working units prior to RoC estimation to allow for uneven temporal distribution of the analysed levels within the sequence using selective binning with a moving window.
3. Estimating compositional dissimilarity between working units.
4. Detecting statistically significant peak-points in the RoC estimates within the sequence.

Given the various choices of coefficients, methods, and evaluations in RoC analysis, it is essential when presenting RoC results for stratigraphical assemblage data to document what choices were made. There are, as we have shown, many decisions in RoC analysis. Publishing plots of ‘Rate of Change’ with no explanation about what choices were made in the RoC analysis should be strongly discouraged (cf. Abrook et al., 2019).

Our RoC analysis approach as implemented in *R-Ratepol* is, we propose, a significant improvement over existing methods for RoC analysis that do not incorporate methods to detect statistically significant peak-points (Birks, 2012). In addition, *R-Ratepol*, used with an appropriate dissimilarity coefficient, can be used to estimate RoC for various types of proxies such as pollen, diatoms, chironomids, cladocerans, molluscs, and sediment chemical variables, among others. Therefore, *R-Ratepol* is, we believe, a powerful and much needed addition to the toolkit of robust numerical techniques available to palaeoecologists and palaeolimnologists for detecting and summarising patterns in complex multivariate time-ordered stratigraphical sequences (Birks, 2010, 1997).

## 5. Acknowledgements

All the authors are supported by the European Research Council under the EU Horizon 2020 Research and Innovation Programme (grant 741413 HOPE) Humans on Planet Earth – Long-term impacts on biosphere dynamics. HJBB is indebted to John Line (University of Cambridge) for collaboration in 1990 in the development of the Fortran77 RATEPOL program, which was a starting point for R-Ratepol, to Cathy Jenks for indispensable editorial help, and to Hilary Birks for many helpful discussions. We are indebted to Gavin Simpson for valuable comments on an earlier version of the manuscript and to Richard Telford for critical evaluation of the underlying methodology. Data were obtained from the Neotoma Paleoecology Database (www.neotomadb.org) and its constituent European Pollen Database (EPD). The valuable work of the data contributors, data stewards, and the Neotoma and EPD community is gratefully acknowledged.

## 6. Authors’ contribution

OM and HJBB conceived the ideas and designed the study. OM led the manuscript writing, data analyses, and interpretation of the results. OM, AWRS, KPB, VAF, and SGAF designed the methodology for extracting and processing palaeoecological sequences from Neotoma. OM, JAG, AWRS, MJS, and HJBB contributed to methodology development. HJBB acquired the funding. All authors contributed to the writing of the text and interpretation of the results and gave final approval for publication.

## 7. Data availability

*R-Ratepol* is available as an R package at GITHUB (https://github.com/HOPE-UIB-BIO/R-Ratepol-package), including all the data used in this study. Scripts for all analyses used in this study can be found at GITHUB repository (https://github.com/HOPE-UIB-BIO/RateOfChange).

