## Supplementary Methods, Tables, and Figures for "Rate-of-change analysis in palaeoecology revisited: a new approach"

Running headline: ROC in palaeoecology revisited

Supplementary Material

Detailed description of individual steps for computation of RoC in R-Ratepol, see also Fig. 1:

1. Assemblage and age-model data are extracted from the original source and should be compiled together, i.e. depth, age, variable (taxon) 1, variable (taxon) 2, etc.

2. (Optional) Smoothing of assemblage data: Each variable within the assemblage data is smoothed using one of five in-built smoothing methods:
   a. None: data are not smoothed
   b. Shepard's 5-term filter (Davis, 1986; Wilkinson, 2005): Smoothing over 5 points following the equation: $\mathrm{V}_{NEW} = \frac{17*\mathrm{V} + 12*(\mathrm{V}_{(+1)} + \mathrm{V}_{(-1)}) - 3*(\mathrm{V}_{(+2)} + \mathrm{V}_{(-2)})}{35}$ , where V is the focal level value. All result values smaller than zero are treated as zero.
   c. Moving average (Wilkinson, 2005): Each value is calculated as the average over $N$ number of levels (preferably ½ $N$ before and ½ after the selected level; values are adjusted at the beginning and end of the sequence. $N$ must be an odd number). Note that each calculation is done from scratch and results are saved separately in order to avoid cumulative rounding errors. Default $N$ is set to 5.

d. Age-weighted average (Wilkinson, 2005): Similar to the moving average but the average is weighted by the age difference from the focus level and multiplied by $\frac{1}{\mathrm{CONSTANT}}$ (CONSTANT is selected by the user). To avoid up-weighting levels, if $\frac{\mathrm{CONSTANT}}{AGEDIFF}$ exceeds 1, it is treated as 1. This means that levels closer than the CONSTANT to the target age are given full weight, but those farther away are downweighted by an amount increasing with age difference. CONSTANT has a default setting of 500 years.

e. Grimm's smoothing (Grimm & Jacobson, 1992): Similar to a moving average but $N$ is not fixed. For each level, $N$ is selected as an odd number between $N_a$ and $N_b$ while maintaining the maximum age difference from the selected levels as $Age_{MAX}$. Default values are set as $N_a$= 5, $N_b$ = 9, $Age_{MAX}$ = 500.

3. Creation of time bins: As the size and position of time bins is pre-defined, a template for all time bins in all window movements can be created.
   RoC can be calculated between individual levels or bins. If binning or binning with a moving window is selected as the working unit (WU), bins of selected size are created as a template, starting from the beginning of the sequence. If binning with a moving window is selected, the creation of bins is repeated many times, with each bin (window) shifting by Z years forward. This is repeated X times, where X = bin size / Z.

4. Single run (an individual loop) is computed:
   a. (Optional) Selection of one time series from the age uncertainties: One age sequence is randomly selected from the uncertainties generated by the age-depth models. Each of the levels is given one age value and the correct temporal order is maintained in the sequence.
   b. Subsetting levels in each bin: If levels are selected as WUs, each level becomes one WU. If bins are selected, levels in predefined bins are pooled together and one level will be selected from each time bin as a representative. The default setting is that levels closest to the beginning of the time bin are selected, but an optional setting is that one level is selected randomly. All other levels are excluded. Bins with valid levels become the new WUs, and empty bins are excluded. The age of each time bin is selected as middle value of its age limits.
   c. (optional) Standardisation of assemblage data in each WU: Each WU is standardised to a selected number of individuals (e.g. 150 pollen grains), using random sampling of individuals without replacement. The dataset is reduced by excluding all rows and columns (WUs and taxa) lacking any data.

d. Calculate RoC between WUs: First, the dissimilarity coefficient between WUs is calculated using one of five dissimilarity coefficients (Gower, 1971; Prentice, 1980; Legendre & Legendre, 2012):

i. Euclidean distance:

$\sqrt{\sum_{i=1}^{m}(A_i - B_i)^2}$, where $A_i$ and $B_i$ are variable values for WUs A and B of variable $i$, and there are *m* variables. This is appropriate when the absolute amounts of the variables are critical, such as pollen influx or concentration data.

ii. Standardised Euclidean distance:

$\sqrt{\sum_{i=1}^{m}(\frac{A_i - B_i}{SD_i})^2}$ , where $A_i$ and $B_i$ are variable values for WUs A and B of variable $i, SD_i$ is the standard deviation for variable $i$ calculated from the whole sequence, and there are *m* variables. This is appropriate when the variables are expressed in different units, as in sedimentary, chemical, physical, or biogeochemical data.

iii. Chord distance:

$\sqrt{\sum_{i=1}^{m}(\sqrt{A_i} - \sqrt{B_i})^2}$, where $A_i$ and $B_i$ are relative variable values for WUs A and B of variable $i$, and there are *m* variables (Prentice, 1980). This and the next coefficient are appropriate with closed compositional data containing many zero values.

iv. Chi-squared coefficient:

$\sqrt{\sum_{i=1}^{m}\frac{(A_i - B_i)^2}{(A_i + B_i)}}$, where $A_i$ and $B_i$ are relative variable values for WUs A and B of variable $i$ , and there are *m* variables (Prentice, 1980). This is the most appropriate coefficient to use with closed (percentage or proportional) compositional data containing many zero values (e.g. pollen percentage data).

v. Gower's (1971) distance:

$1 - \left(\frac{1}{p}\sum_{i=1}^{p} s_i(A_i, B_i)\right)$, where $s_i(A_i, B_i)$ is a partial similarity function computed separately for each descriptor. Therefore, it can be used with data with missing values and assemblages containing mixed data types (e.g. quantitative, qualitative, binary) such as plant macrofossil data (Birks, 2014).

a. Next, the age difference between subsequent WUs is calculated. The RoC score between WUs is calculated as: $RoC_{A-B} = \frac{DC_{A-B}}{Age.diff_{A-B}}$ , where $DC_{A-B}$ is the dissimilarity between WUs A and B, and $Age.diff_{A-B}$ is the age difference between WUs A and B. In addition, the RoC between WUs can be calculated using every consecutive WU, or alternatively, the calculation of RoC can be restricted to only directly adjacent WUs. Using the former increases the number of samples for which RoC can be calculated within a sequence, which varies in terms of sample resolution, but it may still introduce biases related to the RoC estimation as a result of the varying inter-sample distances.

   e. Summarisation of results from all moving windows: If binning with a moving window is selected for the selection of WUs, bins (windows) are moved forward by a selected amount of time $Z$, and the steps 4a–4d are repeated for a new set of WUs (age sampling, levels selection, RoC calculation). This is repeated $X$ times, where $X = \frac{BIN\ size}{Z}$. Results from all window positions are merged and saved as a single value.

5. Step 4 is repeated multiple times, with a default number of 10,000 randomisations.

6. Validation and summary of results from all randomisations.
Results from all randomisations are merged while keeping the identity of each WU combination. Due to random selection of the age sequence, each of the WU combinations will result in multiple RoC values. Final RoC values are calculated as the median score of all randomisations. In addition, due to excluding empty bins, there is a chance that some WU combinations will be present only in some randomisations (if age uncertainties are being used). Therefore, only WU combinations that are present in at least 10% of all randomisations are retained.

7. Data beyond the selected age are excluded (e.g. 8000 cal yr BP for the pollen sequences in this study).

8. Detection and validation of significant peak-points
A rapid change in composition or relative abundances of taxa within the sequence can provide a means of comparing RoC between sequences and interpreting the potential drivers of assemblage change. To detect such significant peak-points of RoC scores in each sequence, each point is tested to see if it represents a significant increase in RoC values. There are various ways to detect significant peak-points in a time series and R-Ratepol is able detect such peak-points using five methods:

   a. Threshold: Each point in the RoC sequence is compared to a median of all RoC scores from the whole (i.e. *threshold value*). The point is considered significant if the 95$^{th}$ quantile of the RoC scores from all calculations is higher than the *threshold value*.

b. Linear trend: A linear model is fitted between RoC values and their ages. Differences between the model and each point is calculated (residuals). Standard deviation (SD) is calculated from all the residuals. A peak is considered significant if it is 1.5 SD higher than the model (this value can be selected by user).

c. Non-linear trend: A conservative generalised additive model (GAM) is fitted through the RoC scores and their ages ($\mathrm{GAM} = \mathrm{RoC} \sim \mathrm{s}(\mathrm{age}, k = 3)$) using the *mgcv* package (Wood, 2011). The $k$ is fixed to a low value to obtain up to a second polynomial relationship. The distance between each point and the fitted value is calculated (residuals). Standard deviation (SD) is calculated from all the residuals. A peak is considered significant if it is 1.5 SD higher than the model (this value can be selected by user).

d. F-deriv GAM: A smooth GAM model is fitted to the RoC scores and their ages ($\mathrm{GAM} = \mathrm{RoC} \sim \mathrm{s}(\mathrm{age})$. The first derivative as well as continuous confidence intervals are calculated from the model using the *gratia* package (Simpson, 2019). A peak is considered significant if the confidence intervals of the first derivative is higher than 0 (for more information see Simpson, 2018).

e. Signal-to-noise index (SNI) method: We adapted SNI from Kelly *et al.* (2011), which was developed to detect changes in charcoal stratigraphical records. SNI is calculated for the whole RoC sequence and a peak is considered significant if it has an SNI value higher than 3.

### Supplementary Tables and Figures

**Table S1. AIC results in terms of successful detection of peak-points comparing different WU selection methods**. We fitted $\mathrm{glmmTMB}_{success-method}$ with the ratio of WU marked as peak-points to all WUs in the focal area ($R_{success-method}$) as the dependent variable with a beta error distribution. Independent variables were: WU (3 level factor), peak-point detection method (5 level factor), dataset type (4 level factor), and all their interactions. Individual dataset ID and RoC setting (factor combining smoothing method and DC; 10 levels) were selected as the random factors. We then used the *dredge* function from the *MuMIn* package to fit models with all possible combinations of predictors (with a constraint that dataset type must be included) and ordered the models by parsimony (AICc). We only selected the best model if ΔAIC < 2. Plus-symbol (+) indicates that the predictor is included in the model.

- SEPARATE FILE -

**Table S2. AIC results in terms of false positives of peak-points comparing different WU selection methods**. We fitted $\mathrm{glmmTMB}_{FalsePositive-method}$ with the ratio of levels marked as peak-points to all levels at the time of the manual edit of the environmental data ($R_{FalsePositives-method}$) as the dependent variable with a beta error distribution. Independent variables were: WU (3 level factor), peak-point detection method (5 level factor), dataset type (4 level factor), and all their interactions. Individual dataset ID and RoC setting (factor combining smoothing method and DC; 10 levels) were selected as the random factors. We then used the *dredge* function from the *MuMIn* package to fit models with all possible combinations of predictors (with a constraint that dataset type must be included) and ordered the models by parsimony (AICc). We only selected the best model if ΔAIC < 2. Plus-symbol (+) indicates that the predictor is included in the model.

- SEPARATE FILE -

**Table S3. AIC results in terms of successful detection of peak-points using binning with the moving window method and GAM peak-point detection methods**. We fitted $\mathrm{glmmTMB}_{success-detail}$ with $R_{success-detail}$ as the dependent variable with a beta error distribution and the independent variables were data-smoothing type (5 level factor), dissimilarity coefficient (2 level factor), position of environmental change (i.e. density of levels; 2 level factor), richness of dataset (2 level factor), and all their interactions. Individual dataset ID was selected as the random factor. Similarly, the *dredge* and *compare_performance* functions were used to select the best mode and reduce unnecessary predictors. Plus-symbol (+) indicates that the predictor is included in the model.

- SEPARATE FILE -

**Table S4. AIC results in terms of false positives of peak-points using binning with the moving window method and the GAM peak-point detection methods**. We fitted $\mathrm{glmmTMB}_{FalsePosite-detail}$ with $R_{FalsePositve-detail}$ as the dependent variable with a beta error distribution and the independent variables were data-smoothing type (5 level factor), dissimilarity coefficient (2 level factor), position of environmental change (i.e. density of levels; 2 level factor), richness of dataset (2 level factor), and all their interactions. Individual dataset ID was selected as the random factor. Similarly, the *dredge* and *compare_performance* functions were used to select the best mode and reduce unnecessary predictors. Plus-symbol (+) indicates that the predictor is included in the model.

- SEPARATE FILE –

**Table S5. Model comparison result for cases from Table S3, which have similar parsimony ΔAIC < 2.** We selected the best model using the *compare_performance* function from the *performance* package (highest Performance Score).

| formula | AIC | BIC | R2_conditional | R2_marginal | ICC | RMSE | Sigma | BF | Performance_Score |
|---|---|---|---|---|---|---|---|---|---|
| success ~ DC + position + smooth + position:smooth + (1 \| dataset_ID) | -11718.1 | -11636.2 | 0.9722 | 0.6244 | 0.9260 | 0.0462 | 94.605 | 0.0328 | 0.5652 |
| success ~ position + smooth + position:smooth + (1 \| dataset_ID) | -11718.6 | -11643.1 | 0.9722 | 0.6243 | 0.9260 | 0.0462 | 94.567 | 1 | 0.5312 |

| | | | | | | | | | |
|---|---|---|---|---|---|---|---|---|---|
| success ~<br>diversity +<br>position +<br>smooth +<br>position:smooth +<br>(1 \| dataset_ID) | -11717 | -11635.2 | 0.9722 | 0.6247 | 0.9259 | 0.0462 | 94.568 | 0.0191 | 0.2940 |

AIC = Aaike information criterion; BIC = Bayesian information criterion; ICC = interclass correlation coefficient; RMSE = root mean square of error; Sigma = residual standard deviation; BF = Bayes factor. Performance Score ranges from 0 to 1, higher values indicating better model performance. Calculation is based on normalizing all indices (i.e. rescaling them to a range from 0 to 1), and taking the mean value of all indices for each model.

**Table S6. Model comparison results for cases from Table S4, which have similar parsimony ΔAIC < 2.** We selected the best model using the *compare_performance* function from the *performance* package (highest Performance Score).

| formula | AIC | BIC | R2_conditional | R2_marginal | ICC | RMSE | Sigma | BF | Performance_Score |
|---|---|---|---|---|---|---|---|---|---|
| success ~<br>DC +<br>position +<br>smooth +<br>DC:position +<br>position:smooth + DC:position:smooth +<br>(1 \| dataset_ID) | -119346 | -119207 | 0.9682 | 0.9677 | 0.0145 | 0.0069 | 536.4842 | 4.04E-16 | 0.5714 |
| success ~<br>DC +<br>position +<br>smooth +<br>position:smooth +<br>(1 \| dataset_ID) | -119360 | -119278 | 0.9681 | 0.9676 | 0.0145 | 0.0070 | 535.532 | 1 | 0.4435 |
| success ~<br>DC +<br>diversity +<br>position +<br>smooth +<br>position:smooth +<br>(1 \| dataset_ID) | -119358 | -119270 | 0.9681 | 0.9676 | 0.0145 | 0.0070 | 535.5399 | 0.016 | 0.3947 |

AIC = Aaike information criterion; BIC = Bayesian information criterion; ICC = interclass correlation coefficient; RMSE = root mean square of error; Sigma = residual standard deviation; BF = Bayes factor.

Performance Score ranges from 0 to 1, higher values indicating better model performance. Calculation is based on normalizing all indices (i.e. rescaling them to a range from 0 to 1), and taking the mean value of all indices for each model.

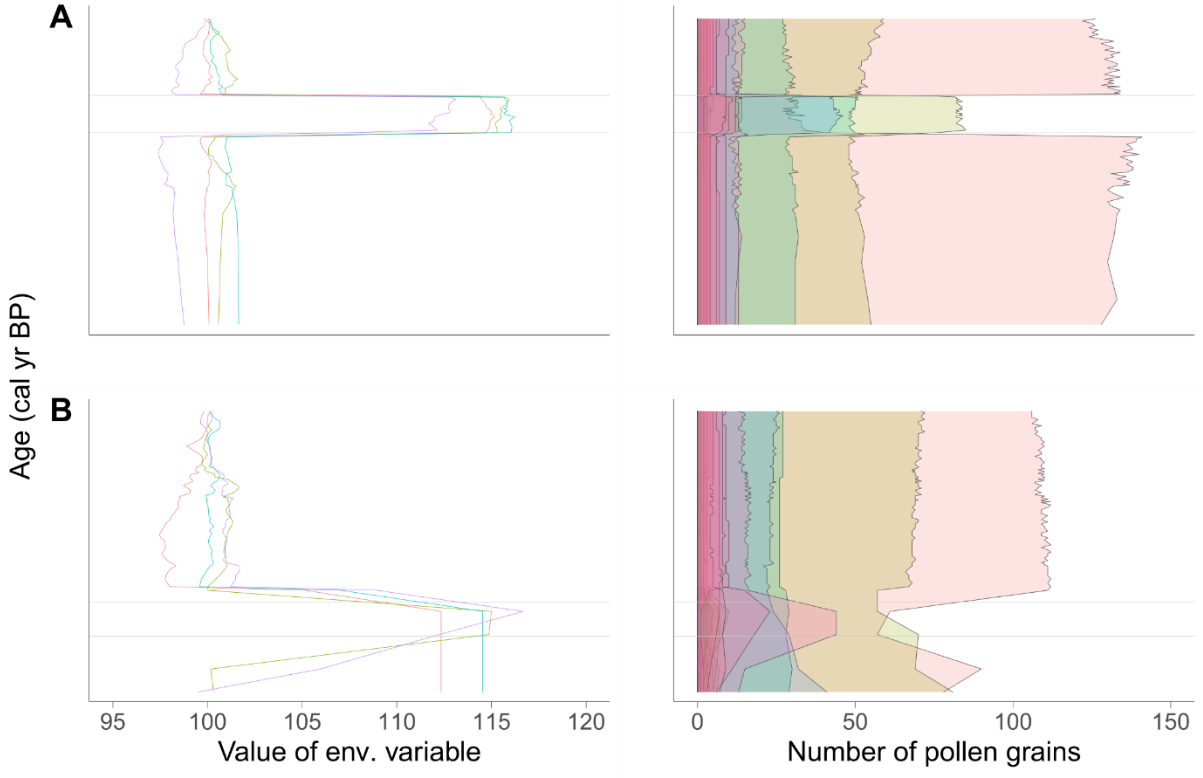


**Figure S1. Example of the manipulation of environmental variables and simulated pollen taxa** in a *late* (A; 3000–2000 cal yr BP) and *early* (B; 6500–5500 cal yr BP) time period, i.e. ‘focal period‘. Note that the *early* focal period has a low density of levels and the *late* focal period has a high density of levels. Left-hand plots show examples of four simulated environmental variables and the right-hand plots show the pollen simulated according to the environmental variables. Changes in environmental variables are reflected in the change of pollen abundances (number of counted pollen grains) of different taxa (right-hand panel different colours)

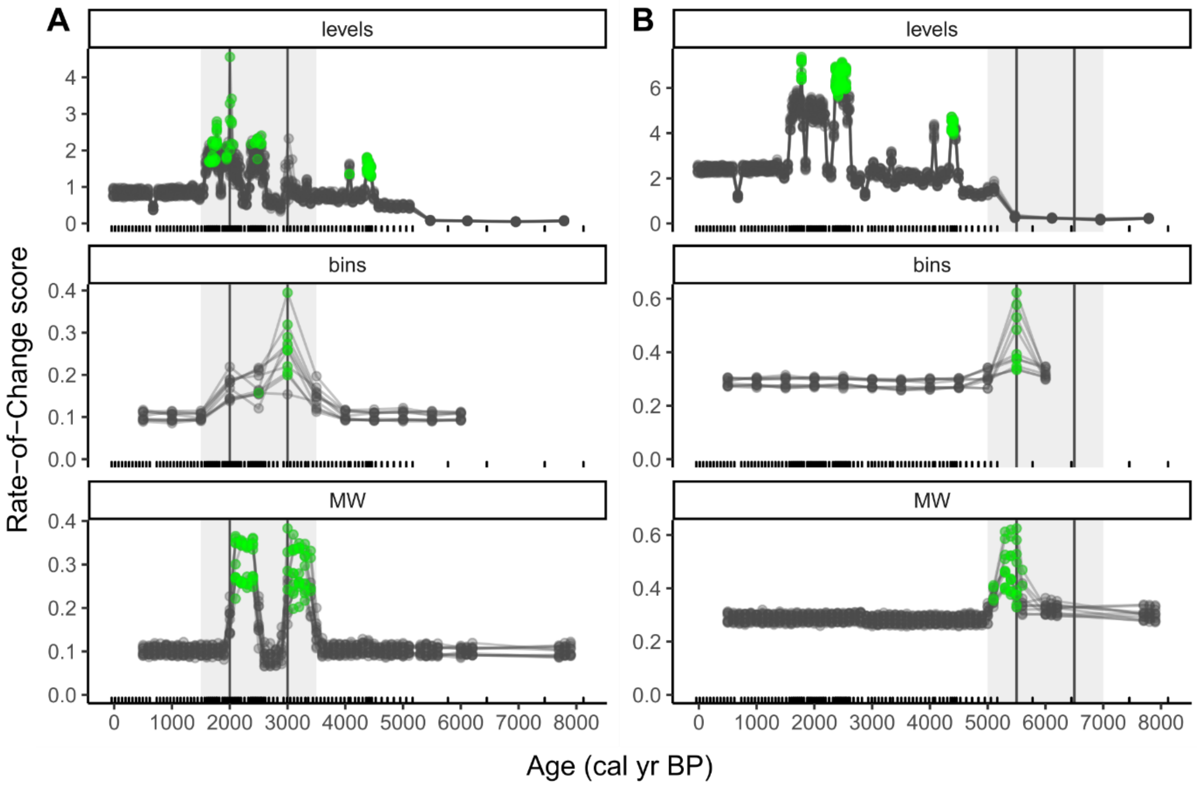


**Figure S2. Example focal periods for success in peak-point detection using simulated pollen data,** estimated using one of the WU-selection methods in a *late* (A) and *early* (B) time period, i.e. 'focal period'. Displayed are all calculations for one example dataset-type (see Methods), with the following properties: richness – high, smoothing method – none, dissimilarity coefficient – Chi-squared, WU specifics – selective binning and binning with a moving window using 500 yr time bins, binning with a moving window using 5 window shifts. RoC score is represented as dissimilarity per 500 yr. Working unit selection: levels = use of subsequent levels (top), bins = selective binning (middle), MW = binning with a moving window (bottom). Each individual line represents one RoC sequence, each point represents one WU RoC estimate. Green points indicate significant peak-points. Vertical lines represent the timings of the change in environmental properties (3000 and 2000 cal yr BP for the *late* period; 6500 and 5500 cal yr BP for the *early* period). The grey background represents the time period used for detecting success.

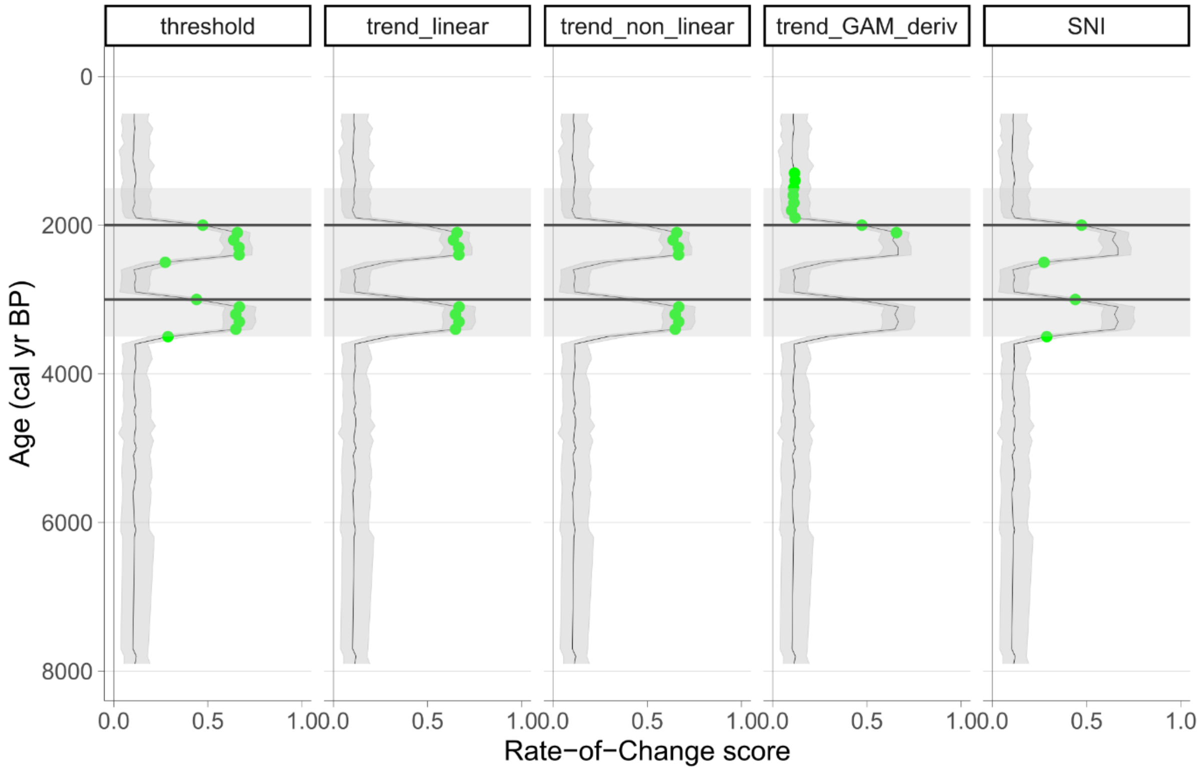


**Figure S3. Comparison of five different methods to detect peak-points in a simulated dataset.** The example dataset has the following properties: richness – high, position of environmental change – *late* (high density of levels), smoothing method – none, dissimilarity coefficient – Chi-squared, Working Unit selection – binning with a moving window with 500 yr bin and 5 window shifts. Horizontal lines represent the timings of the change in the environmental properties (3000 and 2000 cal yr BP for the late period; 6500 and 5500 cal yr BP for the early period). Green points indicate significant peak-points. The grey background represents the time period used for detecting success of the RoC score is represented as dissimilarity per 500 yr. Trend_GAM_deriv = first derivative of a GAM. SNI = signal-to-noise index.

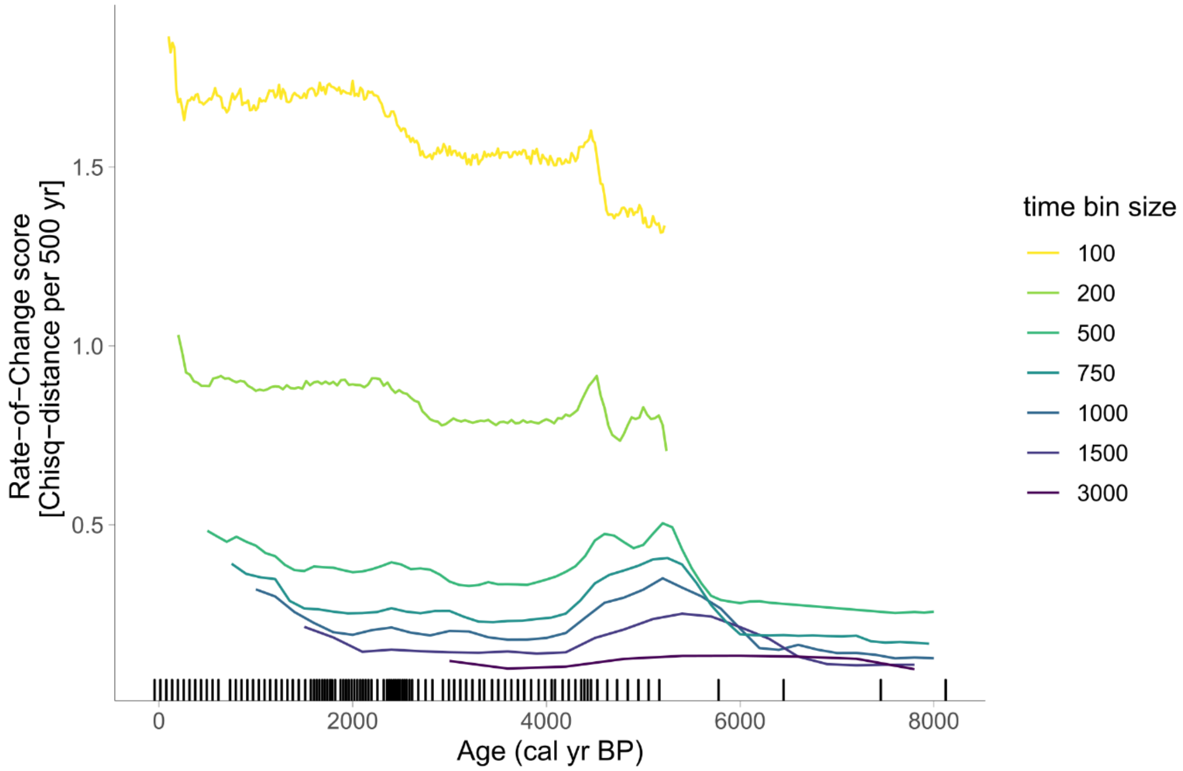


**Figure S4. Rate-of-change scores in Sequence A (Glendalough) calculated with various sizes of time bins using binning with a moving window method.** Age-weighted average and Chi-squared coefficient were selected for smoothing and dissimilarity, respectively. Black ticks on the x-axis indicate the positions of the stratigraphical levels. The RoC score is represented as dissimilarity per 500 yr.

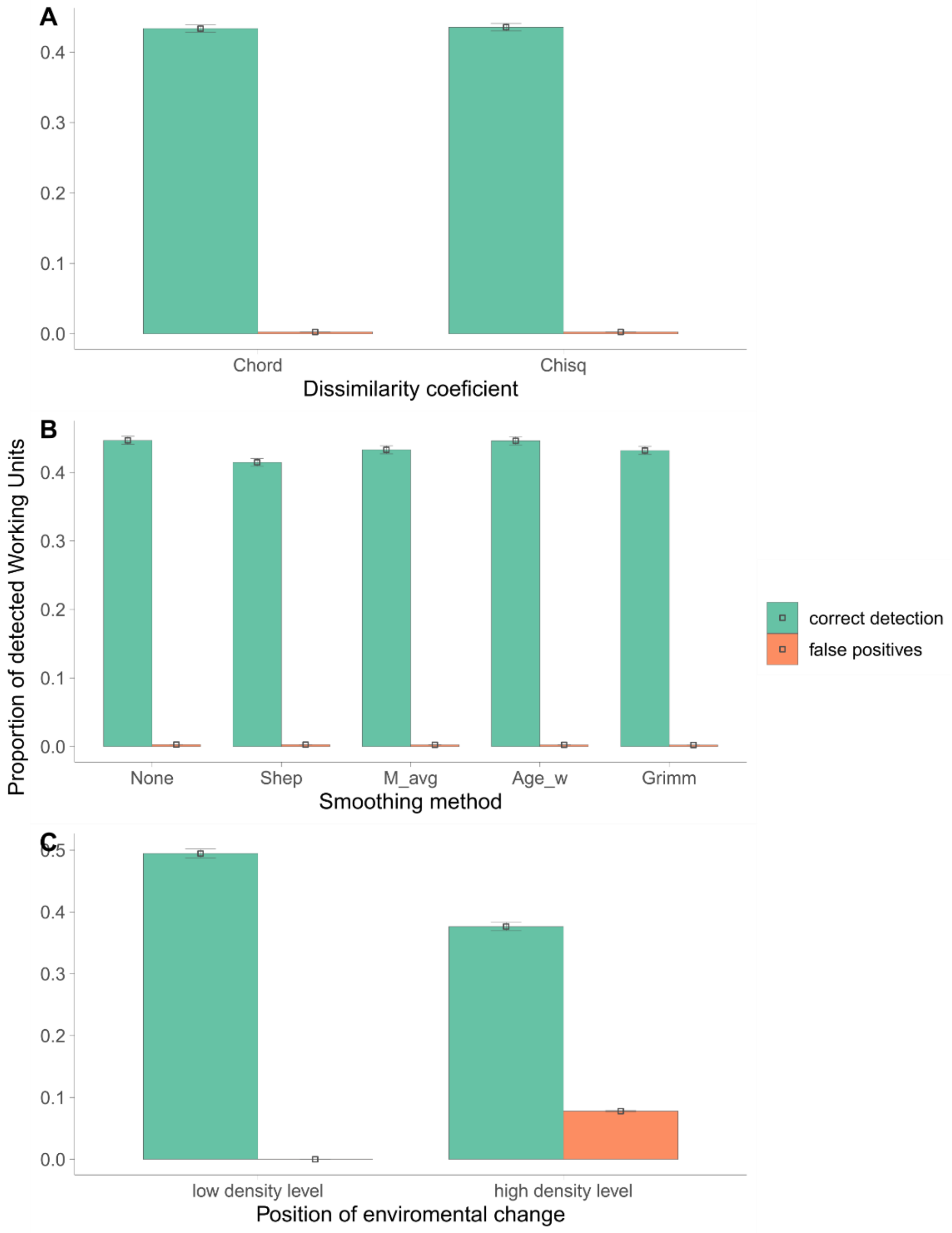


**Figure S5. Comparison of estimated marginal means and 95$^{th}$ quantile of peak-point detection in the simulated datasets for (A) dissimilarity coefficient, (B) smoothing of the data, and (C) density of levels in the period of environmental change**. Dissimilarity coefficient: Chord = Chord distance, Chisq = Chi-squared coefficient. Smoothing of pollen data: None = data without smoothing, Shep = Shepard's 5-term filter, M_avg = moving average, Age_w = age-weighted average, Grimm = Grimm's smoothing. Note that the low density of levels is in the *early focal area* and the high density of levels is in the *late* focal area.

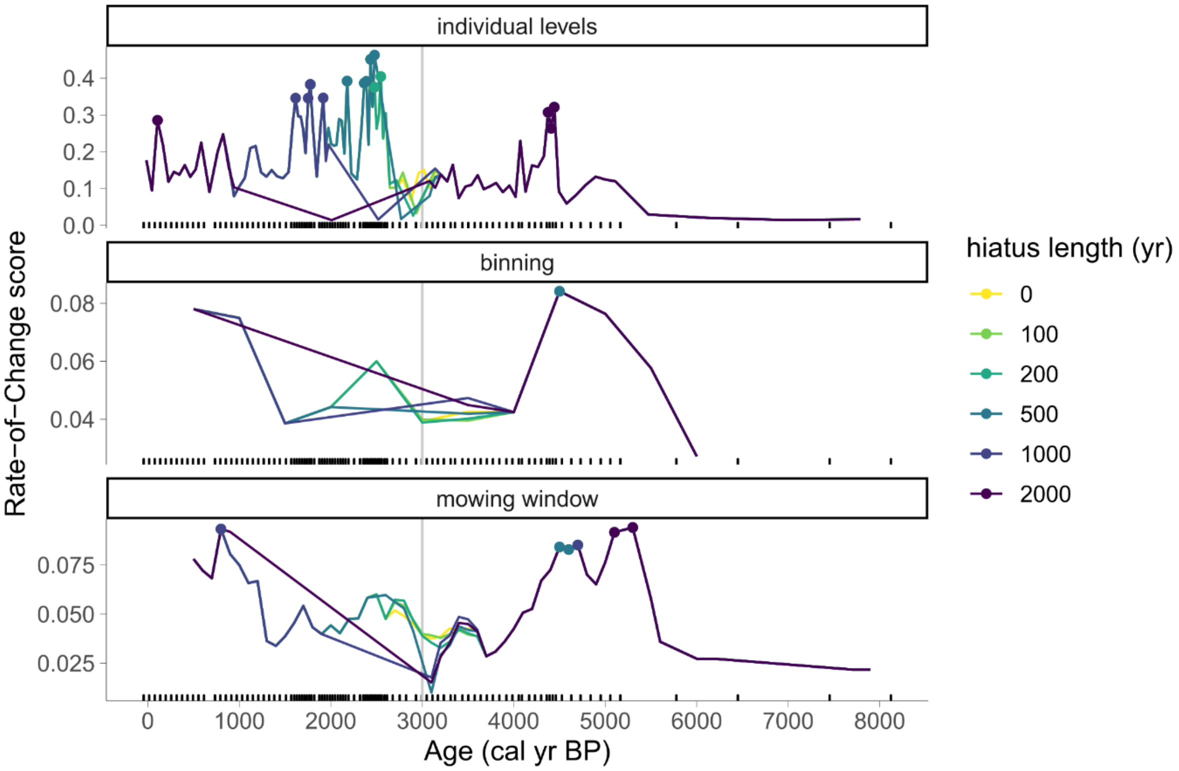


**Figure S6. Sensitivity analyses showing RoC scores for datasets with various hiatus lengths, estimated using three different WU selection methods (individual stratigraphical levels, selective binning, and binning with a moving window)**. A series of datasets with manually created hiatuses of various lengths were created from Sequence A (Glendalough). Age-weighted average and Chi-squared coefficient were selected for smoothing and dissimilarity, respectively. The time bin for binning and binning with a moving window was set as 500 yr. Black ticks on the x-axis indicate the positions of the stratigraphical levels (full dataset). The RoC scores are represented as dissimilarity per 500 yr. The vertical grey line represents the start of the hiatus. Note that the high rates of change detected in RoC in the individual-level approach between 2000 and 3000 years (top panel) are likely to be the result of the changing sedimentation rates in the core. The binning approaches effectively smooth these effects.

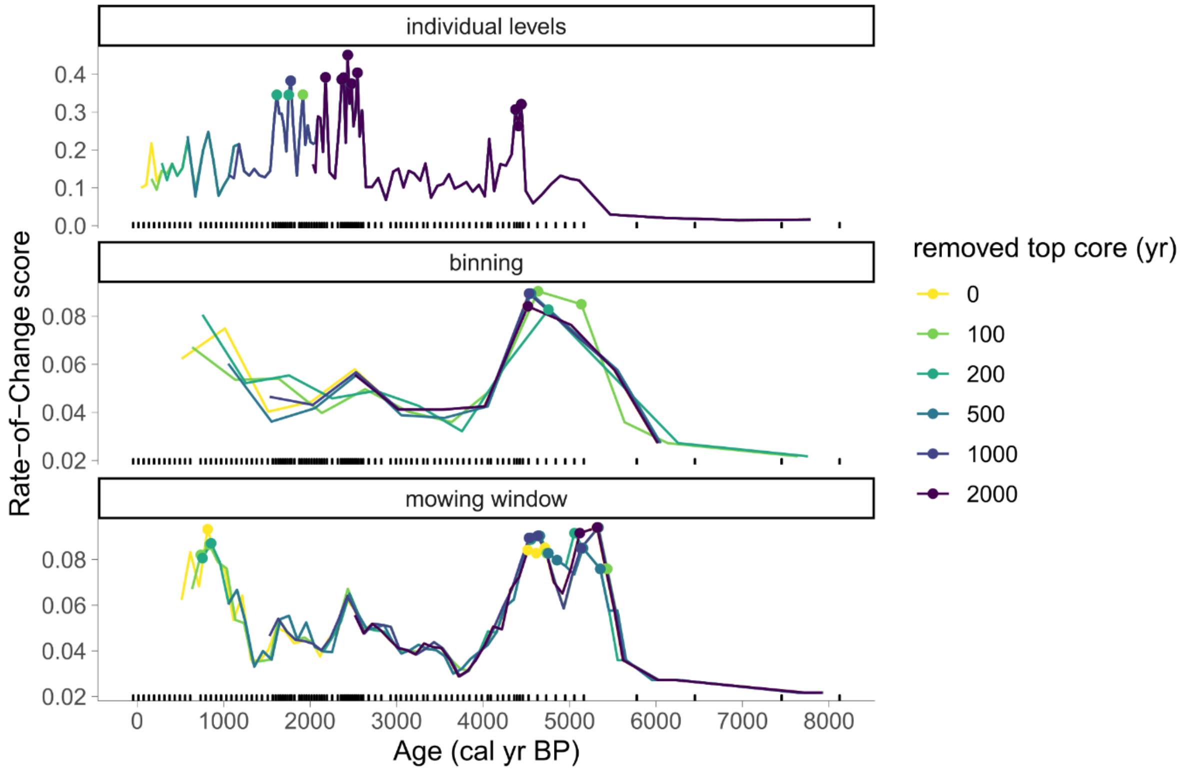


**Figure S7. Sensitivity analyses showing RoC scores for datasets with various hiatus lengths starting from the beginning of the sequence, estimated using three different WU selection methods (individual stratigraphical levels, selective binning, and binning with a moving window)**. A series of datasets, with samples removed starting from the beginning of the sequence for various periods of time, were created from Sequence A (Glendalough). Age-weighted average and Chi-squared coefficient were selected for smoothing and dissimilarity, respectively. The time bin for binning and binning with a moving window was set as 500 yr. Black ticks on the x-axis indicate the positions of the stratigraphical levels (full dataset). The RoC scores are represented as dissimilarity per 500 yr. Note that the high rates of change detected in RoC in the individual-level approach between 2000 and 3000 years (top panel) are likely to be the result of the changing sedimentation rates in the core. The binning approaches effectively smooth these effects.

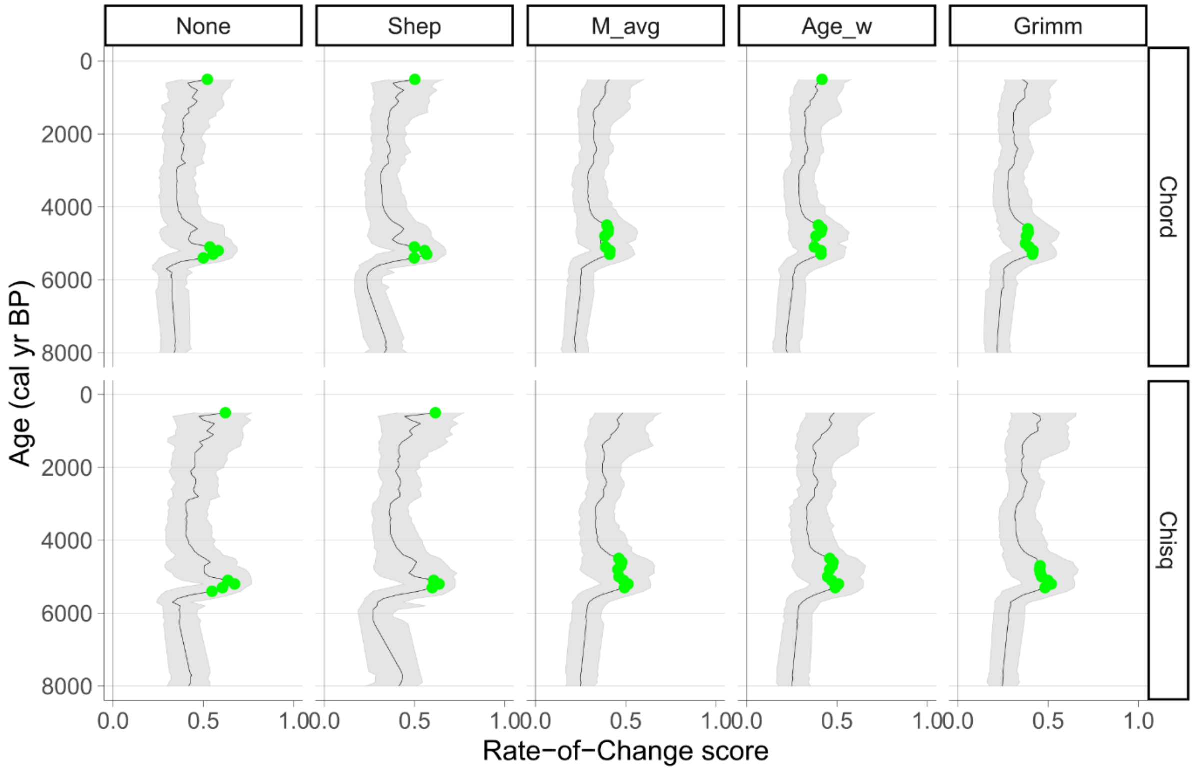


**Figure S8. Rate-of-change (RoC) and peak-points detected in Sequence A (Glendalough) for five categories of data smoothing (columns) on two dissimilarity coefficients (rows).** The binning with moving window was selected as Working Units, with 500 yr bin and 5 window shifts. Green points indicate significant peak-points detected using the non-linear trend method. The RoC scores are represented as dissimilarity per 500 yr. Smoothing methods: None = data without smoothing, Shep = Shepard's 5-term filter, M_avg = moving average, Age_w = age-weighted average, Grimm = Grimm’s smoothing. Dissimilarity coefficients: Chord = Chord distance, Chisq = Chi-squared coefficient. For a detailed explanation of the methods, see Supplementary Material above.
